# Different fluorescent labels report distinct components of spHCN channel voltage sensor movement

**DOI:** 10.1101/2024.01.23.576936

**Authors:** Magdalena N Wojciechowski, Chaseley E McKenzie, Andrew Hung, Alibek Kuanyshbek, Ming S Soh, Christopher A Reid, Ian C Forster

**Affiliations:** Florey Institute of Neuroscience and Mental Health, 30 Royal Parade, Parkville VIC 3052, Australia; School of Science, STEM College RMIT University, GPO Box 2476, Melbourne VIC 3001, Australia; Universität Münster, Institut für Pharmazeutische und Medizinische Chemie, Pharmacampus, Corrensstraße 48, 48149 Münster, Germany

## Abstract

Voltage clamp fluorometry was used to probe the S4 helix movement in the voltage sensing domain of the sea urchin HCN channel expressed in *Xenopus* oocytes. Markedly different fluorescence responses were obtained with either ALEXA-488 or MTS-TAMRA covalently linked to Cys332 at the N-terminal end of S4. With hyperpolarizing steps, ALEXA-488 fluorescence increased rapidly showing characteristics consistent with it reporting the initial inward movement of S4 in agreement with previous studies. In contrast, MTS-TAMRA fluorescence was slower and correlated with the early phase of channel opening. In addition, a slow fluorescence component was resolved with both labels that tracked the development of the mode shift or channel hysteresis. This was quantitated as an increased deactivation tail current delay with concomitantly longer activation periods and was found to depend strongly on the presence of K^+^ ions in the pore. This indicated that the microenvironment of the fluorescent probes attached to Cys332 was strongly influenced by conformational changes in the pore domain. Collisional quenching experiments established that ALEXA-488 was more exposed to solvent than MTS-TAMRA. This was supported by structural predictions based on homology modelling of spHCN in the closed and open conformations with covalently linked fluorophores. This study demonstrates that components of S4 movement during channel activation can be kinetically resolved using different fluorescent probes to reveal three distinct biophysical properties: voltage-sensor movement, early channel opening and mode-shift. These data support the use of different labelling probes to interrogate distinct biophysical aspects of voltage-gated membrane proteins.

**Summary:** Voltage clamp fluorometry was used to probe the S4 helix movement in the voltage sensing domain of the spHCN channel expressed in *Xenopus* oocytes, labeled with either ALEXA-488 or MTS-TAMRA. Each fluorophore reported different components of S4 movement.

## Introduction

Hyperpolarization-activated, cyclic nucleotide-gated (HCN) channels are responsible for cardiac rhythmicity and neuronal excitability (for review see (Biel et al., 2009, Wahl-Schott and Biel, 2009)). They are permeable to Na^+^ and K^+^ ions and are modulated by intracellular cAMP. Unlike other members of the superfamily of voltage-gated potassium (K_v_) channels, HCN channels open at hyperpolarizing potentials, a property that is crucial for their physiological roles. They also differ structurally from other K_v_ channels in that the tetrameric subunit assembly does not involve domain swapping of the transmembrane domains (Flynn and Zagotta, 2018). Our current understanding of HCN channel voltage dependent activation comes from several decades of studies using electrophysiological and fluorometric techniques as well as more recent structural modelling predictions. Channel opening, induced by a rapid hyperpolarization of the membrane potential is initiated by an inward fast movement of the S4 helix of the voltage sensor domain (VSD) (Dai et al., 2019), followed by the development of a helix break in S4 that permits re-orientation of the S5 and S6 domains to allow pore opening through rotation of S6 (Elbahnsi et al., 2022, Wu et al., 2021, Burtscher et al., 2023, Kasimova et al., 2019, Lee and MacKinnon, 2019). Voltage clamp fluorometry (VCF) (for review see (Cowgill and Chanda, 2019, Priest and Bezanilla, 2015) has played a key role in many of these studies by allowing real-time assaying of localised conformational changes associated with voltage sensor movement (Bruening-Wright et al., 2007, Bruening-Wright and Larsson, 2007, Ramentol et al., 2020, Wu et al., 2023, Wu et al., 2021). In its conventional implementation, a cysteine substituted at a chosen site of interest is covalently labeled with a fluorophore and changes in the fluorophore’s microenvironment caused by direct or indirect conformational changes are reported as changes in fluorescence intensity resulting from collisional quenching or other mechanisms (e.g. (Cha and Bezanilla, 1998, Papp et al., 2022)).

Here we characterise the behavior of two commonly used fluorophores (ALEXA-488 and MTS-TAMRA) under the same measurement conditions using excitation and emission hardware appropriate for each label, and show that they report on distinct aspects of HCN voltage-dependent activation. We used the sea urchin sperm HCN channel isoform (spHCN) (Gauss et al., 1998) with an engineered cysteine at position Arg332 at the N-terminal end of the S4 helix (Bruening-Wright et al., 2007) and expressed in *Xenopus* oocytes. ALEXA-488 labeling at this and other sites has been used extensively to study the role of S4 movement in HCN channel gating (Ramentol et al., 2020, Wu et al., 2021, Wu et al., 2023, Bruening-Wright and Larsson, 2007, Bruening-Wright et al., 2007). In contrast, MTS-TAMRA (and related rhodamine dyes), which have been used for VCF studies on electrogenic transporters (e.g. (Patti et al., 2016, Gorraitz et al., 2017, Loo et al., 2005, Virkki et al., 2006)), K_v_ channels (e.g. (Cha and Bezanilla, 1998, Mannuzzu and Isacoff, 2000)) and receptors (e.g. (Papp et al., 2022, Fryatt and Evans, 2014, Dekel et al., 2012)), has so far not been reported as a probe for VCF studies on spHCN channels. In this study, we show that in response to membrane hyperpolarization, ALEXA-488 fluorescence was consistent with a rapid movement of the S4 domain that preceded channel opening in agreement with previous studies (Bruening-Wright et al., 2007, Ramentol et al., 2020, Wu et al., 2023, Wu et al., 2021). In contrast, under the same experimental conditions, the fluorescence emitted by MTS-TAMRA was delayed and correlated with the early phase of channel opening. Both labels also reported a slower component of fluorescence that we show is associated with the mode shift induced by previous hyperpolarization and which depends on the availability of K^+^ ions in the pore. Our findings demonstrate that the use of different fluorescent labels provides new insights into voltage activation of HCN channels and will serve as a valuable tool to study drug interactions with HCN channels.

## Materials and methods

### Solutions and reagents

Defolliculated oocytes were washed in standard OR-2 solution that contained (in mM) 82.5 NaCl, 2 KCl, 1 MgCl_2_.6H_2_O, 5 HEPES, adjusted to pH 7.4 with TRIS. Injected *Xenopus laevis* oocytes were incubated in a standard ND96 storage solution that contained (in mM): 96 NaCl, 2 KCl, 1 MgCl_2_.6H_2_O, 1.8 CaCl_2_.2H_2_O, 5 HEPES, adjusted to pH 7.4 with TRIS and supplemented with antibiotic gentamicin (50mg/l). Electrophysiology was performed using a superfusing solution that contained (in mM) 100 KCl or 100 NaCl, 1.8 BaCl_2_, 1 MgCl_2_, 10 HEPES, pH 7.4 adjusted with TRIS (100K or 100Na solution). Ba^2+^ was substituted for Ca^2+^ to minimise contamination from endogenous Cl^-^ and K^+^ channel currents. CsCl was added as powder to the 100K working solution to give a final concentration of 10 mM. For iodide quenching experiments, powdered KI was added to a working solution in which 50% of KCl was replaced with choline chloride to give 50 mM I^-^ (Patti et al., 2016). The higher osmolarity of this solution was well tolerated by the oocytes.

### cRNA preparation

cDNA encoding the wild type spHCN (spHCN^WT^) was kindly supplied by Dr R Seifert (MPINB, Bonn) and cDNA for the mutant spHCN-R332C (spHCN^R332C^) channel was kindly supplied by Prof H-P Larsson (University of Miami, Miami). To improve the protein expression in oocytes, the cDNA was subcloned into the pGEMHE-MCS vector containing the 5’ and 3’ UTRs from *Xenopus* β-globin. The correct DNA sequence was verified by Sanger sequencing. The plasmid was linearized with *Nhe*I-HF (New England Biolabs) and purified using QIAquick PCR Purification Kit (QIAGEN). The quality and concentration of the linearized plasmid were confirmed *via* gel-electrophoresis and Nanodrop Spectrophotometer (Thermo Fisher Scientific). Following in *vitro* transcription to cRNA was performed using the linearized cDNA template and the mMessage mMachine® T7 transcription kit (Ambion, Thermo Fisher Scientific, Waltham, MA). For cRNA purification the RNeasy Mini Kit (QIAGEN) was used. RNA concentration and purity was verified using NanoDrop Spectrophotometer. cRNAs were stored at -80°C.

### Expression in Xenopus laevis oocytes

Adult female *Xenopus laevis* frogs were anesthetized with 1.3 mg/ml tricaine methanesulfonate (MS-222) and ovaries surgically removed via a small abdominal incision in accordance with animal handling protocols. Oocytes were defolliculated with 1.5 mg/ml collagenase for 1 h and washed with calcium-free OR-2 solution. Mature oocytes stage V or VI were selected and injected manually with 10 ng of cRNA in a 50 nl injection solution, and were maintained in ND96 storage solution at 17 °C for 2-3 days before use.

### Two electrode voltage clamp

Standard two-electrode voltage clamp (TEVC) hardware was used (TEC-05X, NPI. Tamm, Germany), with series resistance (R_s_) compensation applied to ensure clamp accuracy when oocytes exhibited high functional expression. Micropipettes were filled with 3 M KCl with resistances between 0.2-0.5 MΩ. Voltage clamp control and data acquisition were under software control (pClamp version 10, Molecular Devices). All experiments were performed at 18-20 °C. Oocytes were placed in a small perfusion chamber (filled volume ∼20 μl) that was continuously perfused by gravity feed via a common manifold and flow was controlled by pinch valves. Solution exchange occurred in typically < 4 s (McKenzie et al., 2023). Data were sampled at 50 µs/point and low pass filtered at 1000 Hz using the TEC-05X in-built analog Bessel filter.

### Voltage clamp fluorometry

The voltage clamp fluorometry (VCF) apparatus comprised the TEVC (see above) and a laboratory-built fluorescence microscope mounted below the recording chamber cover slip as previously described (Virkki et al 2006). For experiments using MTS-TAMRA labeling, oocytes were incubated in the dark at room temperature for approximately 15 min in a 100K solution supplemented with 40 μM of the label. For experiments using ALEXA-488, oocytes were incubated for 30-60 min in a 100K solution supplemented with 100 μM of the label. Following incubation, oocytes were washed in 100K solution and stored in the dark on ice in 100K solution to reduce internalization of labeled channels. For oocytes labeled with ALEXA-488, changes in fluorescence emission were resolved using a XF100-2 cube set (475AF40 excitation filter, 505DRLP dichroic mirror and 535AF45 emission filter) (Omega Optical Inc) and excited by a blue LED (L1RX-BLU) (Luxeon, Philips LumiLEDs). For oocytes labeled with MTS-TAMRA, changes in fluorescence emission were resolved using a XF32 cube set (535DF35 excitation filter, 570DRLP dichroic mirror and 596DF35 emission filter) (Omega Optical Inc) and excited by a green LED (L1RX-GRN), (Luxeon, Philips LumiLEDs). The LEDs were driven by a stabilized, current source (typically 300-500 mA, depending on the fluorophore) and cell exposure to the light source was controlled by an electronic shutter (VS252T1, Vincent Assoc, USA). After labeling, oocytes were placed animal pole down in the perfusion chamber. The fluorescence signal was detected by a photodiode (S1336-BQ, Hamamatsu) and the diode current converted to a voltage using an Axopatch 200A (Molecular Devices) amplifier. Diode bias was adjusted to minimize noise and headstage resetting. The output was further amplified, low-pass filtered (8 pole Bessel) and sampled at the same rate as for the TEVC signals. Some photobleaching occurred after multiple light exposures and extended protocols before and after ligand application. Scaling corrections were made based on changes in background fluorescence, where we assumed that all labeled proteins were equally affected. Oocytes expressing spHCN^WT^ channels, gave no Δ*F* in response to voltage steps after incubating with MTS-TAMRA under the same conditions indicated above, as previously reported for ALEXA-488 (Bruening-Wright and Larsson, 2007) (data not shown), which established that labeling of native Cys was unlikely to contribute to Δ*F*.

### Data analysis

Data were analysed using Prism 9.1 (Graphpad; USA) and Clampfit 10.4 (Molecular Devices, USA). Time constants for activation were obtained from exponential fits using Clampfit routines applied to a region commencing after the initial inflexion to the end of the test pulse. For strong hyperpolarising potentials, the activation current time course was best described using a double exponential fit (Hung et al., 2021, Bruening-Wright et al., 2007). The fluorescence signal was described by either a double exponential (MTS-TAMRA labeling) or a variable offset rising exponential (ALEXA-488 labeling) as indicated. All exponential fits used the Chebychev fitting algorithm in Clampfit. The macroscopic conductance (*G*), was derived from the instantaneous tail currents, by assuming a constant driving force and number of channels. First, a baseline correction was made after the current had reached a steady-state at +40 mV and the instantaneous current was measured at the start of the repolarising step (*I*_tail_) immediately after the capacitive transient had settled and fit with a form of the Boltzmann equation:

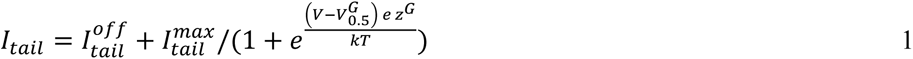

where *I_tail_^off^* is the maximum instantaneous tail current and I_tail_^off^ is the offset reported by the fit (typically =0 with baseline correction), *V*_0.5_*^G^* is the mid-point voltage, *Z^G^* the valence of the apparent charge moved, and *kT*/*e*=25.3 mV at 20 °C. A similar equation was used to characterise the Δ*F-V* data with corresponding parameters: midpoint voltage (*V*_0.5_*^F^*) and valence (*Z^F^*) of the apparent charge moved. Data sets were normalised to the predicted maximum and pooled from replicate data sets. All data points are shown as mean ± SEM and all pooled experimental data were obtained from oocytes from >2 donor frogs unless otherwise noted.

### Kinetic simulations

Kinetic simulations of a 12 state model for HCN channels (Wu et al., 2021) were performed using Berkeley Madonna (v.8.3.9), Berkeley CA.

#### Structural modelling

The wild type sea urchin HCN (spHCN^WT^) (spHCN; (UniProtKB entry Q45ZY2) in depolarized and hyperpolarized state conformations were constructed using the SWISS-MODEL server (Waterhouse et al., 2018) with the human HCN1 PDB structures 5U6O and 6UQF (Lee and MacKinnon, 2017, Lee and MacKinnon, 2019), respectively, as structural templates. Residue Arg332 (spHCN^WT^; equivalent to Arg258 in HCN1) in both structures was mutated to Cys using Discovery Studio Visualizer 2021 (Dassault Systèmes). The structures of unliganded fluorophores ALEXA-488 and MTS-TAMRA were constructed using the small molecule build and geometry optimisation modules of DSV.

Structures of depolarized and hyperpolarized spHCN^R332C^ covalently ligated to ALEXA-488 or MTS-TAMRA via Cys332 were obtained as follows.

First, autodock Vina 1.1.0 (Eberhardt et al., 2021, Trott and Olson, 2010) was used to predict initial (non-covalent) binding positions of these two fluorophores at spHCN^R332C^ to identify binding poses that place the ligating atom of these molecules in proximity to Cys332. Single monomers of depolarized and hyperpolarized spHCN^R332C^ were used for docking calculations, as the sizes of the fluorophores and tethering to Cys332 preclude them from forming multi-subunit interactions. The GUI PyRx 0.8 (Dallakyan and Olson, 2015) was used to prepare the Autodock Vina calculations. Blind docking was performed for both fluorophores at both depolarized and hyperpolarized spHCN^R332C^, with the docking box set to ‘maximise’ in all cases. All rotatable torsions were set to be flexible for the two unligated fluorophores. An exhaustiveness parameter value of 48 was used for all Vina docking calculations. For each fluorophore, we selected the most energetically favored pose in which the molecules are oriented such that their Cys-linking atom was close to Cys332.

Second, DSV was used to manually define a covalent bond between Cys332 and the respective fluorophore via a C-S bond for ALEXA-488, and a disulfide (S-S) bond for MTS-TAMRA to produce depolarized and hyperpolarized spHCN^R332C^ covalently liganded to each fluorophore at position 332. This was followed by energy minimization, using DSV, of the covalently-attached fluorophores and residues Val324 to Cys332. This resulted in minor structural corrections for the depolarized spHCN^R332C^, but partial unwinding of this helical segment for hyperpolarized spHCN^R332C^, as Cys332 is buried within the extracellular pocket of the VSD (S4) helical bundle in the hyperpolarized state and would require at least local structural disruption to accommodate the fluorophores. Third, the covalent docking protocol implemented in AutodockFR (Bianco et al., 2016) was then used to predict energetically favored positions of ALEXA-488 and MTS-TAMRA while taking into account orientational rigidity imposed by covalent linkage to Cys332.

A docking box, with grid spacing of 0.375 Å, centred at Cys332, was defined for both depolarized and hyperpolarized conformations, with dimensions of 20 x 25 x 50 Å^3^ to encompass the entire volume hypothetically accessible to the tethered fluorophores. AutogridFR (Zhang et al., 2019) was used to prepare affinity maps, including intra-receptor gradients. Autodock (Morris et al., 2009) was subsequently performed using100 independent docking runs, each using up to 10^5^ evaluations, for each fluorophore and spHCN^R332C^ conformation. The highest ranked poses for ALEXA-488 and MTS-TAMRA at depolarized and hyperpolarized spHCN^R332C^ were selected for subsequent analysis.

Schrödinger Maestro 12.9 (Release 2021-23, Maestro, Schrödinger) was used to obtained 2D ligand interaction diagrams for the covalent docking-obtained results.

## Results

### Distinct voltage dependent fluorescence changes observed after labeling spHCN^R332C^ channels with ALEXA-488 or MTS-TAMRA

We recorded changes in fluorescence (Δ*F*) from oocytes expressing the spHCN^R332C^ channel that were labeled with either ALEXA-488 or MTS-TAMRA in response to hyperpolarizing voltage steps from a 0 mV holding potential. Recordings from two representative oocytes superfused with 100K solution from the same donor frog revealed that the time course of Δ*F* during channel activation was markedly different for each labeling condition (Fig 1A). In response to the voltage step protocol (inset), ALEXA-488 labeled oocytes displayed an initial rapid rise of Δ*F*, followed by a slower component that reached a steady-state by the end of the 1 s test pulse. The rapid rise of Δ*F*, which was easily resolved for the largest hyperpolarizing steps, was slower than the capacitive current transient (see inset, Fig 1A). The 10-90% rise time of this rapid Δ*F* component for steps from 0 mV to −160 mV was 3.95 ± 0.29 ms (n = 6) and was also not limited by the bandwidth of the fluorescence signal conditioning hardware, which had a rise time of 1.75 ms (400 Hz bandwidth). When the membrane potential was stepped back to +40 mV (for instantaneous current tail analysis (see below)), Δ*F* decayed monotonically and the rapid change in Δ*F* at step onset was absent when returning from the largest hyperpolarizing test potentials. In contrast, MTS-TAMRA labeled oocytes displayed a slower increase in Δ*F* that commenced at completion of the capacitive transient (see inset, Fig 1A) and did not reach a steady-state during the 1 s hyperpolarizing test pulse. The deactivating decay time course was also monotonic and slower than for ALEXA-488 labeled cells. Moreover, the Δ*F* deactivation phase for both labeling conditions was also markedly slower than the deactivation phase of the current tails, which indicated that the channel pore closed well before Δ*F* returned to the baseline. For both current and Δ*F*, the deactivation time course was well described by a single exponential decay with time constants *τ_deact_^I^* and *τ_deact_^F^*. For a step from −160 mV to +40 mV, ALEXA-488 labeling resulted in *τ*_deact_*^I^* = 15.2 ± 1.3 ms (n=5) and *τ*_deact_*^F^* = 87.2 ± 8.8 ms; MTS-TAMRA labeling resulted in *τ^I^* = 68.0 ± 6.6 ms (n=5) and *τ*_deact_*^F^* = 314.4 ± 48.9 ms. For unlabeled cells expressing spHCN^R332C^, *τ*_deact_*^I^* = 24.6 ± 1.8 ms (n=3). This established that although labeling with MTS-TAMRA had slowed the channel closure rate more markedly than ALEXA-488, the ratio of deactivation rates (τ_deact_^F^/τ_deact_^F^) was similar for each label (ALEXA-488: 5.7; MTS-TAMRA: 4.6). This confirmed that both fluorophores reported on molecular rearrangements involving S4 movement after channels had closed. Moreover, it suggested that the molecular coupling between channel closure and change in S4 microenvironment sensed by each fluorophore upon deactivation was similar.

**Figure 1.**
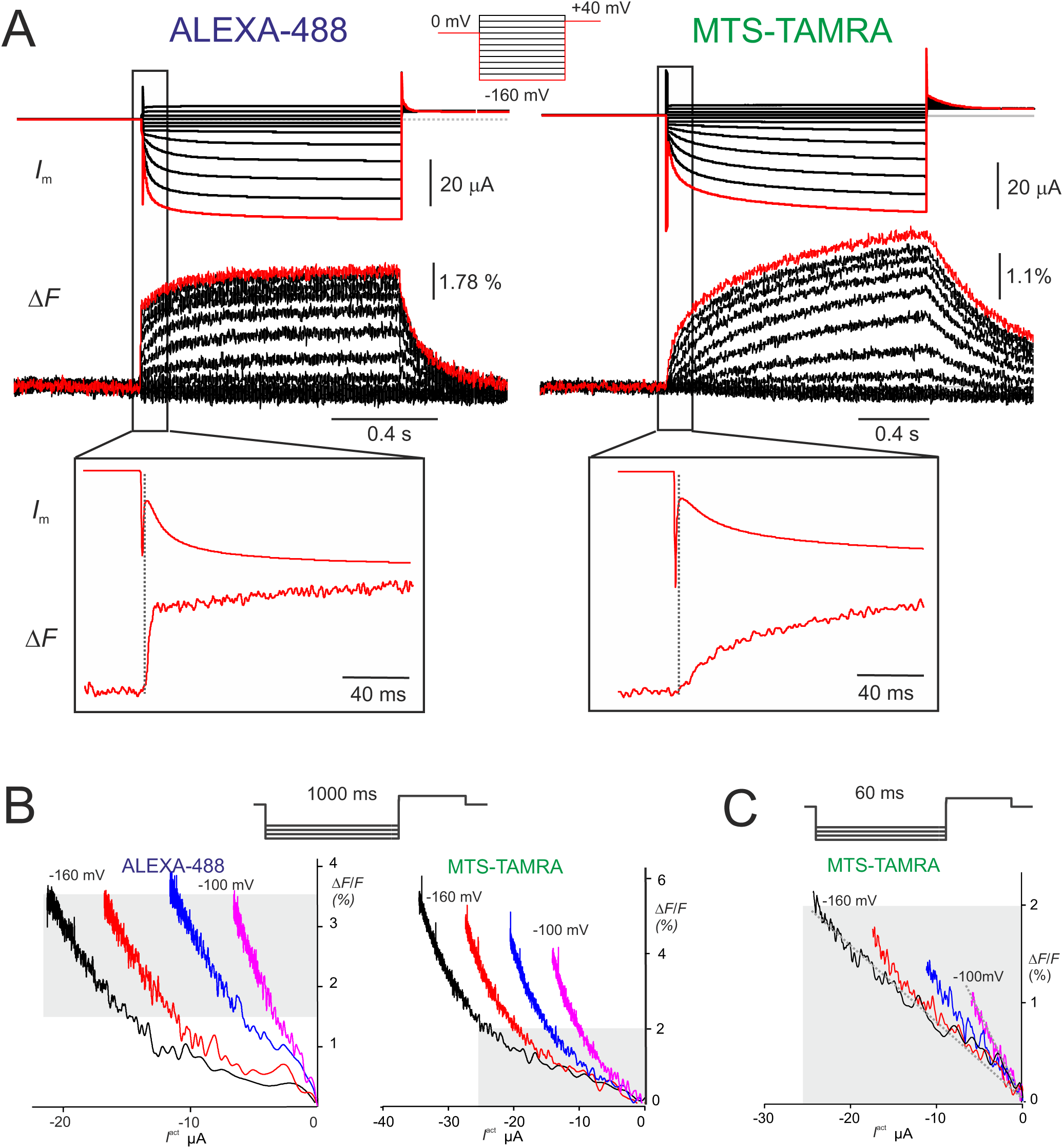
Distinct voltage dependent fluorescence changes by labeling spHCN^R332C^ with ALEXA-488 or MTS-TAMRA. A) Representative recordings of membrane current (*I*_m_: upper panels) and simultaneous fluorescence (Δ*F*-lower panels) for ALEXA-488 labeled oocyte (left) and MTS-TAMRA labeled oocyte (right) in response to the voltage step protocol shown Oocytes were from same donor frog recorded on consecutive days. Boxed regions expanded (x10) for voltage steps to -160 mV highlight the initial fast Δ*F* for ALEXA-488 labeling. The vertical dashed lines indicate the approximate end of the capacitive charging phase. Δ*F* shown as % of background fluorescence. Current and fluorescence were low pass filtered at 400Hz. B) Parametric plots of Δ*F* and activating currents (*I*^act^) for representative oocytes labeled with either ALEXA-488 (left panel) or MTS-TAMRA (right panel) for 4 hyperpolarizing potentials from -100 mV to -160 mV in 20 mV intervals. For the ALEXA-488 data, grey shading indicates region approximately 60 ms after step onset, for which *I*^act^ and Δ*F* appear to correlate. For the MTS-TAMRA data, the initial 60 ms is shaded and expanded in C. C) MTS-TAMRA data for the same oocyte as in B) but using a 60 ms test pulse with same potentials. Grey marked zone corresponds to same area in B) (right panel). Superimposed lines highlight linear relation between early phase of Δ*F* and *I*^act^ for MTS-TAMRA labeling for -100 mV and -160 mV.

Parametric plots at four activation potentials for representative cells indicated that there was no obvious correlation between the overall time course of Δ*F_total_* and activating current (*I*^act^) for either label (Fig 1B). However, for MTS-TAMRA labeling the initial phase of Δ*F* (up to 60 ms) tracked the corresponding rapid initial growth in *I*^act^ (Fig 1C), which suggested that this label reported a conformational change that directly related to channel opening. For times >60 ms after the hyperpolarizing step onset, the parametric plots suggested that Δ*F* and *I*^act^ also tracked for ALEXA-488. Taken together, these findings established that despite being covalently linked to the same cysteine, each label reported different changes to their respective microenvironments as the channels activate and deactivate.

We determined the dependence on membrane potential of the normalised conductance (*G*) from instantaneous tail current analysis (see Materials and Methods) and compared this with the normalised total fluorescence change (Δ*F_tot_*) for each label measured at the end of the test pulse (Fig 2A). For both fluorophores, the midpoint voltage of Δ*F_tot_* (*V*_0.5_*^F^*) was right shifted relative to that of *G* (*V*_0.5_*^G^*) by 43 mV (ALEXA-488) and 38 mV (MTS-TAMRA) (Table 1). This indicated that each fluorophore reported on conformational changes that preceded channel opening, which was in general agreement with previous findings for spHCN^R332C^ labeled with ALEXA-488 (Bruening-Wright et al., 2007). Labeling oocytes expressing spHCN^R332C^ with either fluorophore also altered the voltage dependence of channel activation: for MTS-TAMRA, when compared with the normalized *G-V* data obtained from unlabeled oocytes expressing spHCN^R332C^, *V*_0.5_*^G^* was right shifted by ̴ 38 mV, whereas ALEXA-488 labeling resulted in a smaller right shift of 12 mV (Table 1). Thus, labeling with ALEXA-488 influenced the spHCN^R332C^ channel open probability less than MTS-TAMRA, consistent with the altered deactivation rates for current (see above). We found that spHCN^WT^ expressing oocytes typically showed a −70 mV shift in *V*_0.5_*^G^* compared with spHCN^R332C^ (Table 1) (see also (Mannikko et al., 2002)). The larger right shift of *V*_0.5_*^G^* for MTS-TAMRA suggested that the presence of this label led to a partial compensation for the charge loss that resulted from removal of Arg332 for spHCN^WT^.

**Figure 2.**
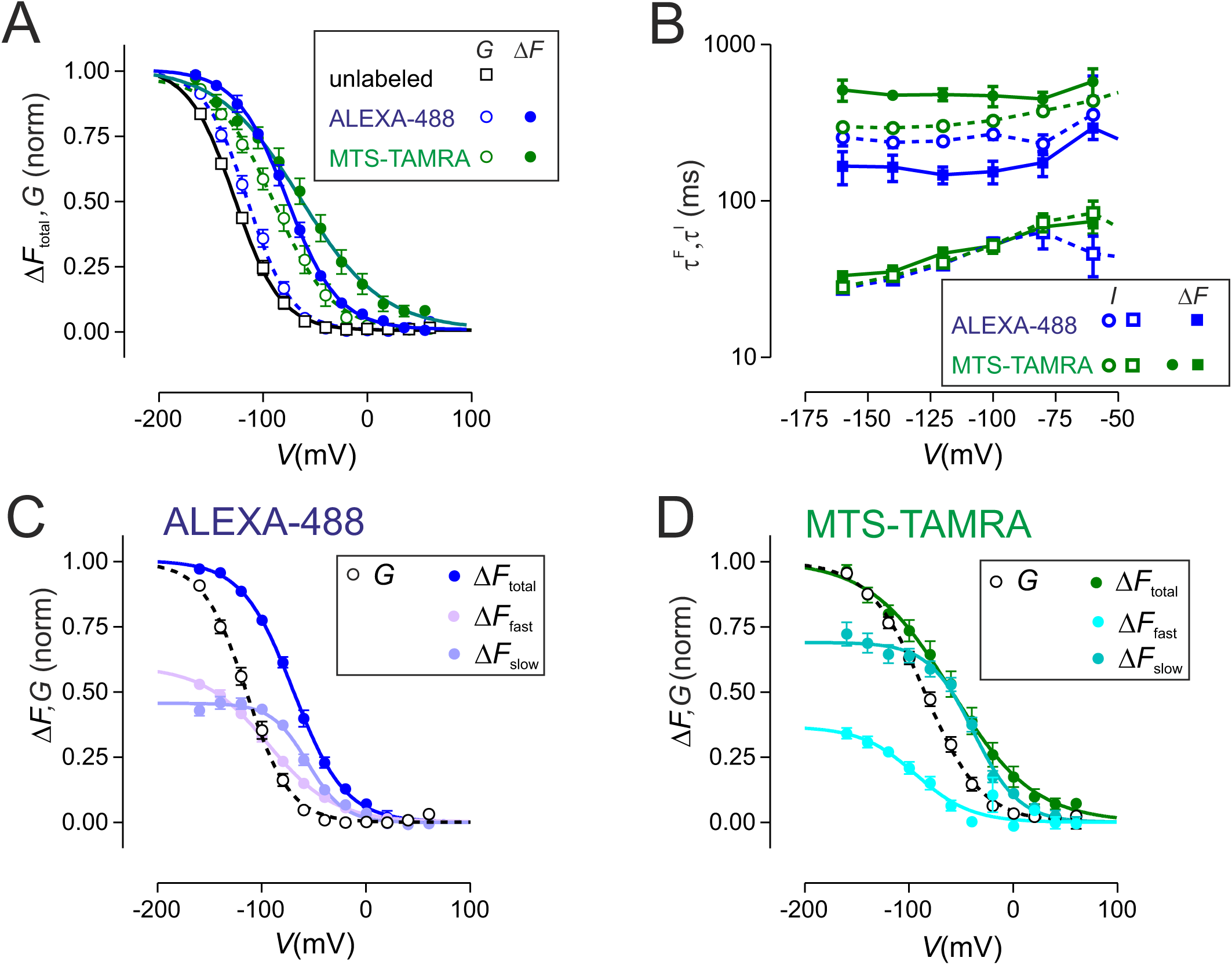
Comparison of voltage dependent properties of unlabeled and labeled spHCN^R332C^. A) Conductance (*G*) derived from tail currents and total Δ*F* (Δ*F_total_*) measured at end of 1 s test pulse normalised to respective maxima predicted from Boltzmann fit to data sets from individual oocytes and then pooled (see Materials and Methods). Open symbols: *G*; filled symbols (Δ*F_total_*. Unlabeled spHCN^R332C^: n=3; ALEX-488 labeled spHCN^R332C^: n=5; MTS-TAMRA labeled; n=5. Symbols shown in boxed inset. B) Activation time constants plotted as a function of test pulse for current (τ^I^) and Δ*F* (τ^F^). For both labels, activation currents were fit with two exponential: for ALEX-488 labeled oocytes, Δ*F* was fit with a single rising exponential with variable offset for ALEXA-488 labeling and a double exponential for MTS-TAMRA labeling. ALEXA-488 labeled spHCN^R332C^: n=15; MTS-TAMRA labeled spHCN^R332C^: n=9. Symbols shown in boxed inset. C) For ALEXA-488 labeled oocytes total Δ*F_total_* was resolved into two components (Δ*F_fast_*, Δ*F_sloss_*) when fit with a single rising exponential with variable offset, where Δ*F_fast_* represents the fit offset and Δ*F_sloss_* represents the amplitude of the exponential component. Same data set as in B). Data shown normalised to Δ*F_total_*. Symbols shown in boxed inset. D) For MTS-TAMRA labeled oocytes Δ*F_total_*was resolved into two components (Δ*F_fast_*, Δ*F_sloss_*) when fit with a double exponential function. Same data set as in B). Data shown normalised to Δ*F_total_*. Symbols shown in boxed inset.

**Table 1.**
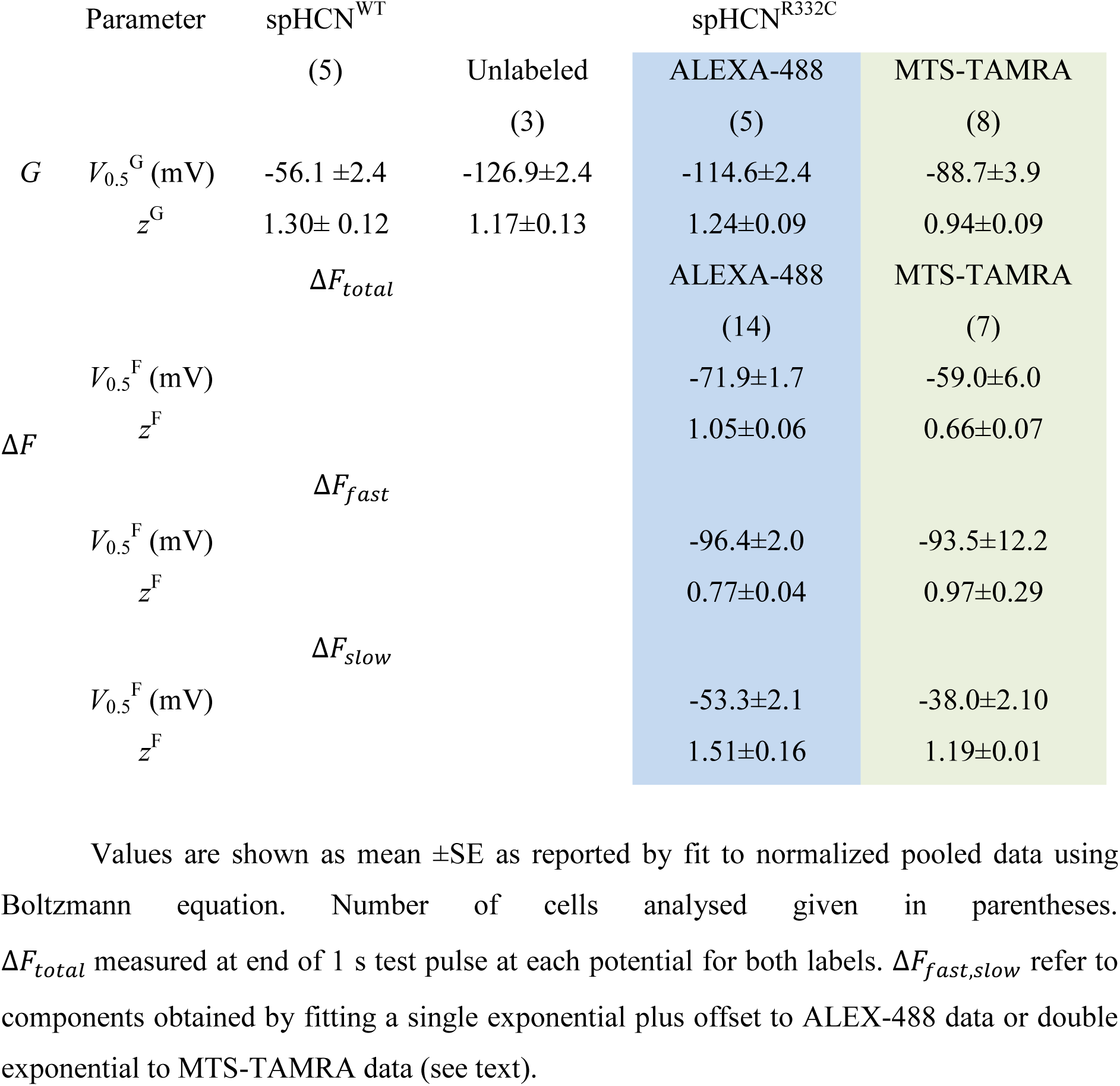
Comparison of Boltzmann fit parameters from normalised conductance (*G*) or change in fluorescence (Δ*F*) for spHCN^WT^, spHCN^R332C^ in 100K solution.

To characterise activation kinetics further, we compared the activation time course before and after labeling of the time-dependent (activating) component of membrane current and fluorescence following the voltage step onset (Fig 2B). For labeled oocytes, current activation was well described by fitting a double exponential function commencing after the capacitive transient had settled, in agreement with previous studies (Bruening-Wright et al., 2007). We only considered time constants reported by the double exponential fit for potentials ≤−60 mV, where an obvious time-dependent activation was observed. For both labels, the faster time constant showed a similar voltage dependence for potentials ≤ −80 mV (Fig 2B), whereas the slower components showed a weak voltage dependence over the same voltage range. For ALEXA-488 labeled oocytes, fitting the complete Δ*F* traces with a standard double exponential function resulted in large fit uncertainty at the step onset and we therefore chose to describe Δ*F* by a single rising exponential with variable offset (see Materials and Methods). This allowed us to separate Δ*F_total_* into two components: an initial pseudo instantaneous change (Δ*F_fast_*) and a slower time-dependent component (Δ*F_sloss_*) described by a single exponential rise to the steady state. The time constant for Δ*F_sloss_* showed little voltage dependence up to −60 mV and it was significantly slower than the main fast time constant for current activation (Fig 2B). Both components of fluorescence showed different steady-state voltage dependencies (Fig 2C) with midpoint voltages lying to the right of the corresponding conductance midpoint (*V*_0.5_*^G^*) (Table 1). This indicated that each component reflected unique molecular conformational changes involving S4 that preceded channel opening. Component Δ*F_sloss_* was resolved at potentials well before most channels would be open and increased in relative amplitude to reach a maximum close to *V*_0.5_*^G^* before slightly decreasing at the largest hyperpolarizing potentials. Component Δ*F_fast_* showed an obvious sigmoidicity that, together with its absence at the “OFF” transition (see above), would argue against it arising from an electrochromic effect in which the fluorophore emission is directly proportional to changes in the transmembrane electric field (Priest and Bezanilla, 2015). The similarity of its midpoint voltage (−96 mV) with that reported for gating charges recorded from spHCN^R332C^ channels (−91 mV, see (Bruening-Wright et al., 2007)) strongly suggested that this component reflected changes in the fluorophore’s microenvironment as the charged S4 domain of the VSD moved inward in response to membrane hyperpolarization (Wu et al., 2021).

The Δ*F* traces for MTS-TAMRA labeling were described by a standard double exponential fit as for the activating current. The time constants associated with each component were generally well separated (>5 fold) although resolution of two time constants was difficult for membrane potentials > −60 mV due to limited signal-to-noise ratio (Fig 1A, right panel). The time constant of the slow component showed no systematic voltage dependence for −160 mV ≤*V*≤−60 mV (Fig 2B). To decrease the fit uncertainty we reduced the number of fit parameters by constraining the slow time constant to the mean of the free fit estimates for voltages from −160 mV to −80 mV for individual oocytes. The fast time constant was similar in voltage dependence and magnitude to the fast component of the activating current (Fig 2B). The voltage dependence of the predicted steady-state fluorescence associated with each component (Δ*F_fast_*, Δ*F_sloss_*), was markedly different (Fig 2D). For small hyperpolarizations, Δ*F_sloss_* accounted for most of the total fluorescence and its midpoint potential (̴ −38 mV) obtained from a Boltzmann fit was significantly right shifted relative to that of the corresponding *G* for MTS-TAMRA labeled spHCN^R332C^ channels, whereas Δ*F_fast_* had a midpoint voltage comparable to *V*_0.5_*^G^* (Table 1). Together with the similarity of the fast time constants for current and fluorescence (Fig 2B), this suggested that Δ*F_fast_* reported conformational changes associated with channel opening and was consistent with the early phase of activation depicted in the parametric plot (Fig 1C).

In summary, the ALEXA-488 Δ*F* was consistent with it reporting the initial activation steps involving independent movement of the VSDs, whereas MTS-TAMRA reported on conformational changes associated with channel opening. In addition, both labels reported a second fluorescence component with slower kinetics that did not correlate with channel opening. The most likely functional correlate for this component would be the mode shift or hysteresis, whereby the voltage dependence of activation depends on the previous history of the channel (Bruening-Wright and Larsson, 2007, Elinder et al., 2006, Mannikko et al., 2005) as investigated below.

### ALEXA-488 and MTS-TAMRA bound to the S4 domain report on different aspects of HCN channel gating

To further characterize the origins of Δ*F*, we reduced the test pulse period to 8 ms so that fewer channels would have time to transition to the open state during hyperpolarization. We chose 8 ms as the shortest pulse period that would allow Δ*F* to be resolved reliably and ensure completion of the capacitive transient using the whole oocyte voltage clamp. In addition, signal averaging (typically from 4 or 8 trials) was employed to improve the Δ*F* signal-to-noise ratio. If Δ*F_fast_* for ALEXA-488 labeling were associated with the initial independent VSD movement, we would expect to see this component return rapidly to the baseline fluorescence upon deactivation. Data from a representative ALEXA-488 labeled oocyte (Fig 3A, left panel) showed the characteristic rapid increase in Δ*F* in response to the hyperpolarization. This was followed by a corresponding rapid decrease in direct response to the voltage step returning to +40 mV, together with a slowly relaxing component that diminished in amplitude with weaker hyperpolarizing potentials. In contrast, for a representative MTS-TAMRA-labeled oocyte recorded under the same conditions (Fig 3A, right panel), resolvable Δ*F* was only observed for strong hyperpolarizing test pulses. This would be expected if MTS-TAMRA fluorescence reported conformational changes associated with channel opening given that most channels remained closed during the test pulse.

**Figure 3.**
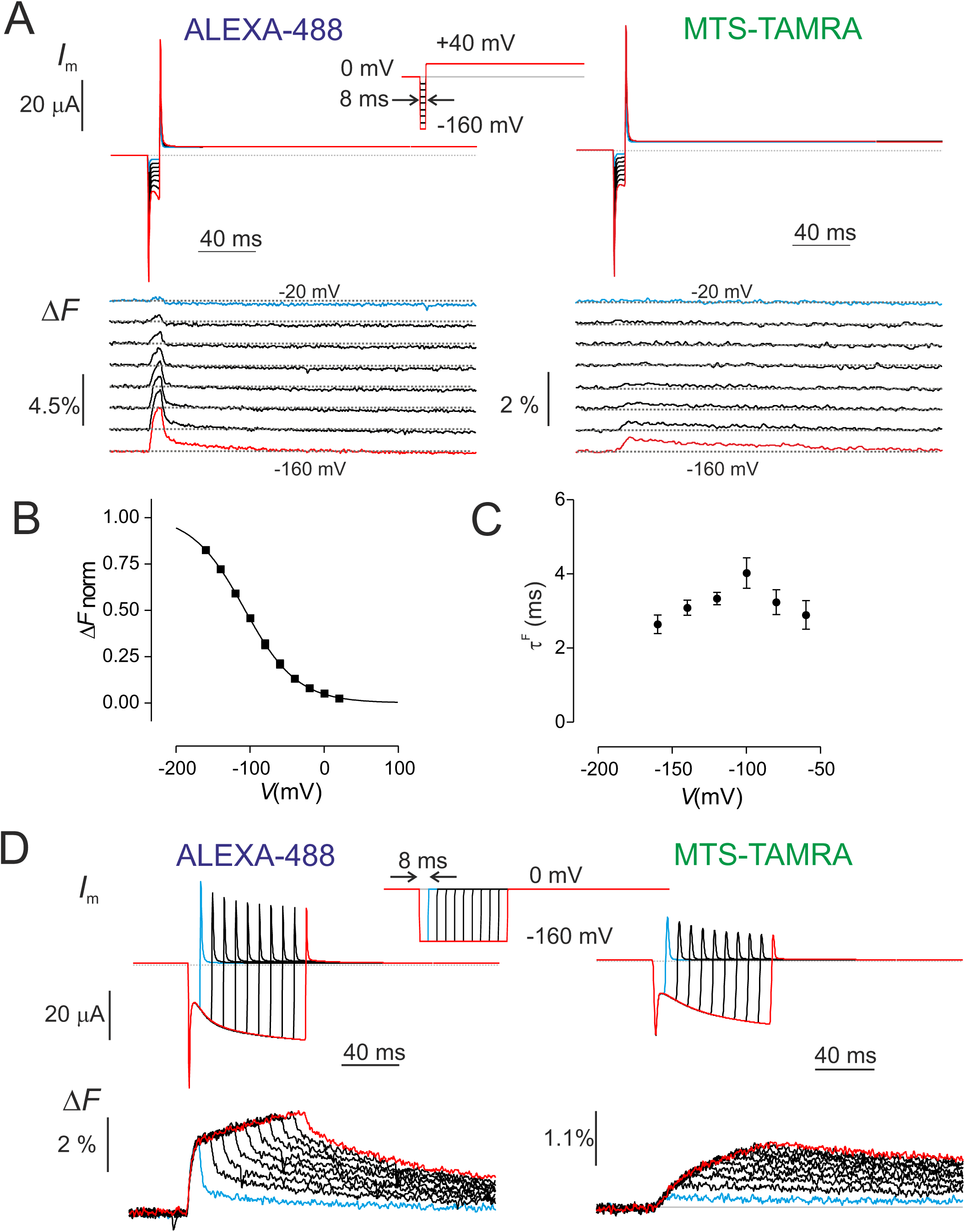
Voltage step protocols highlight different fluorescent responses for each label. A) Representative current and Δ*F* recordings for ALEXA-488 (left) and MTS-TAMRA (right) labeled oocytes using an 8 ms voltage step (inset) showing current (upper) and fluorescence (below). The traces were signal averaged 8-fold to improve the signal-to-noise ratio and separated vertically for visualisation. Δ*F* shown as a % of background fluorescence. B) Δ*F-V* for ALEXA-488 labeled cells. The peak Δ*F* at each voltage at the end of the 8 ms step was first fit with the Boltzmann function and normalised for each cell to Δ*F*^max^ before pooling. Continuous curve is Boltzmann fit with *V*_0.5_^F^= -107.8±5.0 mV; z^F^= 0.73±0.05 ; mean±SE(n=9). C) Voltage dependence of rise-time of Δ*F* for ALEXA-488 labeled cells (same set as in B)) obtained by fitting the rising phase with a single exponential. It was not possible to resolve Δ*F* for voltages above -60 mV. D) Representative current and Δ*F* recordings for ALEXA-488 (left) and MTS-TAMRA (right) labeled oocytes in response to a sequence of prepulses to -160 mV with increasing length (inset) showing current (upper) and fluorescence (below). The traces were signal averaged 8-fold to improve the signal-to-noise ratio. Δ*F* shown as a % of background fluorescence.

For ALEXA-488 labeling, we measured the peak Δ*F* reached at the end of the test pulse. This showed a sigmoidal voltage dependence (Fig 3B) and a Boltzmann fit reported a midpoint voltage (*V*_0.5_*^F^*) of −107.8 ± 5.0 mV and *z*^F^= 0.73 ± 0.05 (n=9). These fit parameters agreed with those for Δ*F_fast_* obtained by fitting Δ*F_total_* obtained from the standard 1 s pulse activation protocol (Table 1) and further supported the notion that this component of Δ*F* reflected the rapid movement of the VSDs. Moreover, a single exponential fit to the rising phase of Δ*F* showed clear voltage dependence (Fig 3C) with a peak close to *V*_0.5_*^F^* that would be expected if both kinetic measures reflected the same voltage dependent transition.

To further characterise Δ*F* for each labeling condition, we used a variable pulse width protocol with 8 ms increments stepping to −160 mV to fully activate channels (Fig 3D). With increasing pulse width, the rapid component of deactivation reported by ALEXA-488 decreased and the amplitude of the slow component of Δ*F* during the deactivating phase grew Fig 3D (left panel). This behavior was consistent with the notion that the activation and deactivation pathways comprise distinct kinetic processes. For the MTS-TAMRA-labeled oocyte, Δ*F* appeared to track the early growth in the activating component of *I*_m_ with increasing pulse width (Fig 3D (right panel), thereby further supporting the previous observation (Fig 1C) that MTS-TAMRA reported conformational changes associated with channel opening, in contrast to ALEXA-488, for which neither component appeared to correlate.

Further insight into identifying the functional correlates of the fluorescent responses was obtained using the 12-state allosteric model proposed recently (Wu et al., 2021) (Fig 4A, see SI for simulation details). As originally proposed, Δ*F* for the ALEXA-488 labeled case comprises two components corresponding to the independent and concerted S4 movements, respectively. In contrast to Wu and colleagues, we assigned a positive intrinsic Δ*F* to both these components to account for our experimental findings. The simulated Δ*F_total_* recapitulated the basic behavior that we observed experimentally if we assume that ALEXA-488 reports both movements, whereas MTS-TAMRA only reports on the slow movement. According to the model, for strong hyperpolarizing potentials, after the initial rise in fluorescence the rapid component (Δ*F*_1_) decreased as more S4 subunits undergo the concerted movement and channels open (Fig 4B). There was a complementary rise in the Δ*F* corresponding to this concerted step (Δ*F*_2_) so that Δ*F_total_* was nearly constant for most of the activation period. The simulations also indicated that there was a close match between the voltage dependence of conductance (*G*) and Δ*F*_2_ as observed experimentally, whereas Δ*F_total_* preceded the channel opening (Fig 4C). Our simulations were also able to recapitulate the variable pulse protocol findings (compare Fig 3D with Fig 4D). However, this model was unable to account for the slower component of Δ*F* observed for both fluorophores, which we now show is related to the mode shift behavior as previously reported (Bruening-Wright and Larsson, 2007).

**Figure 4.**
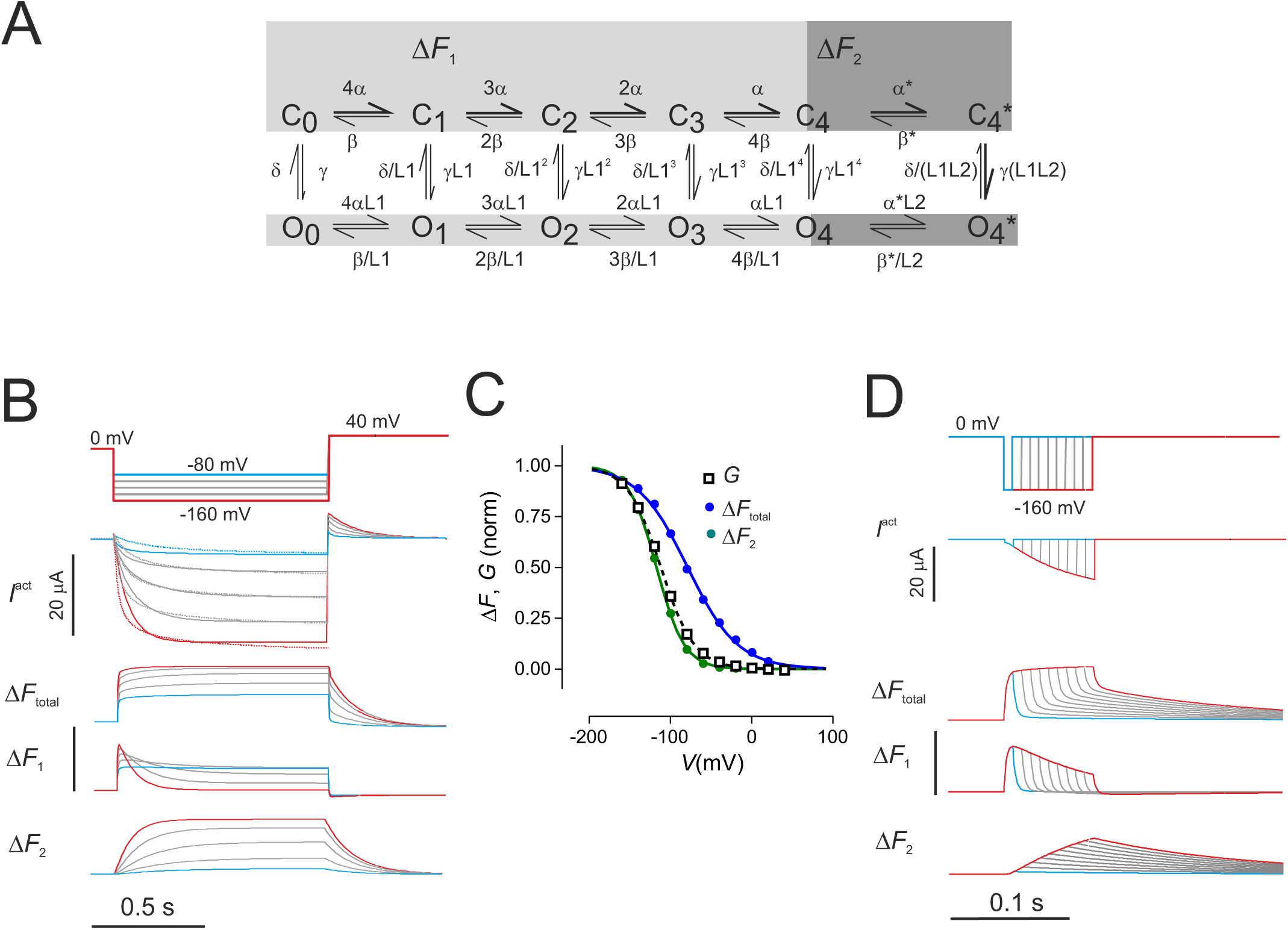
Simulations using an allosteric 12-state model recapitulate features Δ*F* obtained with ALEXA-488 and MTS-TAMRA. A) HCN allosteric model based on the scheme originally proposed by Altomare et al. (Altomare et al., 2001) and extended to 12 states (Wu et al., 2021) was used to simulate activating current and fluorescence using their labeling convention. This scheme does not take account of hysteresis (mode shift), which would require a more complex model with concomitantly more parameters (e.g. see (Mannikko et al., 2005). Vertical transitions represent the closed to open transition of the pore and were assumed electroneutral. Horizontal transitions between states C_0_.._4_ and O_0..4_ are voltage dependent and involve independent movement of S4 helices. If all S4 helices have completed their fast transition, a concerted movement of S4 helices (transitions C_4_-C_4_*, O_4_-O_4_*) is proposed to occur. The forward and backward rates of all voltage dependent transitions were described using the conventional Eyring transition state formulation given by: 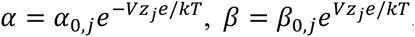, respectively, where *e*, *k* and *T* have their usual meanings, *z* is the apparent valence for the transition. Zero voltage rates α_0_, β_0_ were defined as 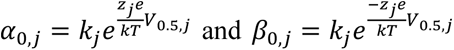, where *j* refers to the fast or slow movement. B) Simulations of *I*^act^, Δ*F* in response to voltage step protocol (top). For simulating *I*^act^, we determined a set of model parameters (Table SI 1) based on a fit to the representative data set of measured activating currents with ALEXA-488 labeling that gave a reasonable match to our data. To simulate the fluorescence in response to voltage steps, we assumed that each of the 12 conformational states can contribute to Δ*F_total_*, i.e. 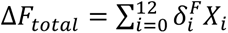, where *X*_i_ is the state occupancy and *δ_i_^F^* is the apparent fluorescence intensity for state i. We assumed that the independent S4 movements (for either open or closed pore) contribute the same fluorescence change per sensor, *δ*_1_*^F^*, 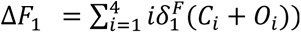, whereas the concerted slow movement contributes a change in fluorescence 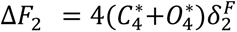. To account for the difference in Δ*F_total_* found experimentally for ALEXA-488 and MTS-TAMRA labeling, we assumed the former reported both fast and slow S4 movements, whereas the latter only reported the slow component. We did not take account of the differences in activation kinetics for the two labels. C) The voltage dependence of normalised conductance (G) and Δ*F* contributions. Δ*F_total_* would correspond to the fluorescence measured by ALEXA-488, whereas Δ*F*_2_ would correspond to that from MTS-TAMRA labeling. Note that the voltage dependence of normalised *G* and Δ*F*_2_ match well. The fit parameters reported by single Boltzmann fit were as follows: *G V*_0.5_=-111.7±0.5 mV, *z*=1.26±0.02; Δ*F_total_ V*_0.5_=-78.9±2.2 mV, *z*=0.84±0.06; Δ*F*_2_ *V*_0.5_=-117.1±0.06, *z*=1.51±0.04 (mean ± SE) D) Simulations of variable prepulse width protocol with an 8 ms increment (compare with Fig 3D). As in B), Δ*F_total_* is proposed to represent the ALEXA-488 labeling case, whereas Δ*F*_2_ would represent the MTS-TAMRA labeling case.

### ALEXA-488 and MTS-TAMRA report development of mode shift in spHCN^R332C^ channels

A previous study of spHCN using fluorescent labeling with ALEXA-488 at sites in S4 other than 332, lying at the N-terminal end of S4, reported a strong correlation between the respective time course of Δ*F* and the temporal development of mode shift (as evidenced by the tail current delay). This observation established that the S4 domain undergoes conformational changes during the mode shift (Bruening-Wright and Larsson, 2007). Moreover, in two earlier studies, the mode shift kinetics for mammalian HCN channels were shown to be strongly associated with the presence of external K^+^ ions in the conductance pathway and their ability to access a binding site within the pore (Mannikko et al., 2005, Elinder et al., 2006). We re-examined these findings in the context of identifying the slow component of Δ*F* observed for each label under our measurement conditions.

We first investigated the effect on Δ*F* of changing the main permeant cations for both labels. The addition of 10 mM CsCl to the standard 100K superfusate (100K+Cs solution), resulted in block of inward activating current and Δ*F_total_^max^* increased by ̴20% (ALEXA-488) and ̴30% (MTS-TAMRA) (Fig 5A, left (ALEXA-488) and right (MTS-TAMRA) panels, Table 2) but only for hyperpolarizing potentials below −60 mV. Substitution of most of the external K^+^ ions with Na^+^ ions (100Na solution containing 2 mM KCl) reduced the activating current as expected for a permeability ratio (P_Na_/P_K_=0.18) (Gauss et al., 1998, Ludwig et al., 1998)) (see insets, Fig 5B). Superfusion with 100K+Cs solution also resulted in increased Δ*F_total_^max^* of 10% for ALEXA488 and 30% for MTS-TAMRA (Fig 5B left (ALEXA-488) and right (MTS-TAMRA), Table 2). The increase in Δ*F_total_* occurred at potentials < −60 mV for both fluorophores, which suggested that they most likely report on the same molecular rearrangements induced by the experimental manipulations that altered the availability of K^+^ ions in the pore (Bruening-Wright and Larsson, 2007). For ALEXA-488 labeling we also found for both solution manipulations that Δ*F_fast_* remained essentially unchanged (Fig 5C, D, upper panels) with no significant shift in *V*_0.5_^F^, and *z*^F^ (Table 2). This established that the rapid S4 movement reported by ALEXA-488 was insensitive to these altered pore permeation conditions or complete block. On the other hand, Δ*F_sloss_* significantly increased in both cases for potentials below −60 mV , peaked at −100 mV and then decreased at the strongest hyperpolarizations (Fig 5C, D, lower panels), reminiscent of the negative Δ*F* component recently reported under comparable superfusion conditions (Wu et al., 2021).

**Figure 5.**
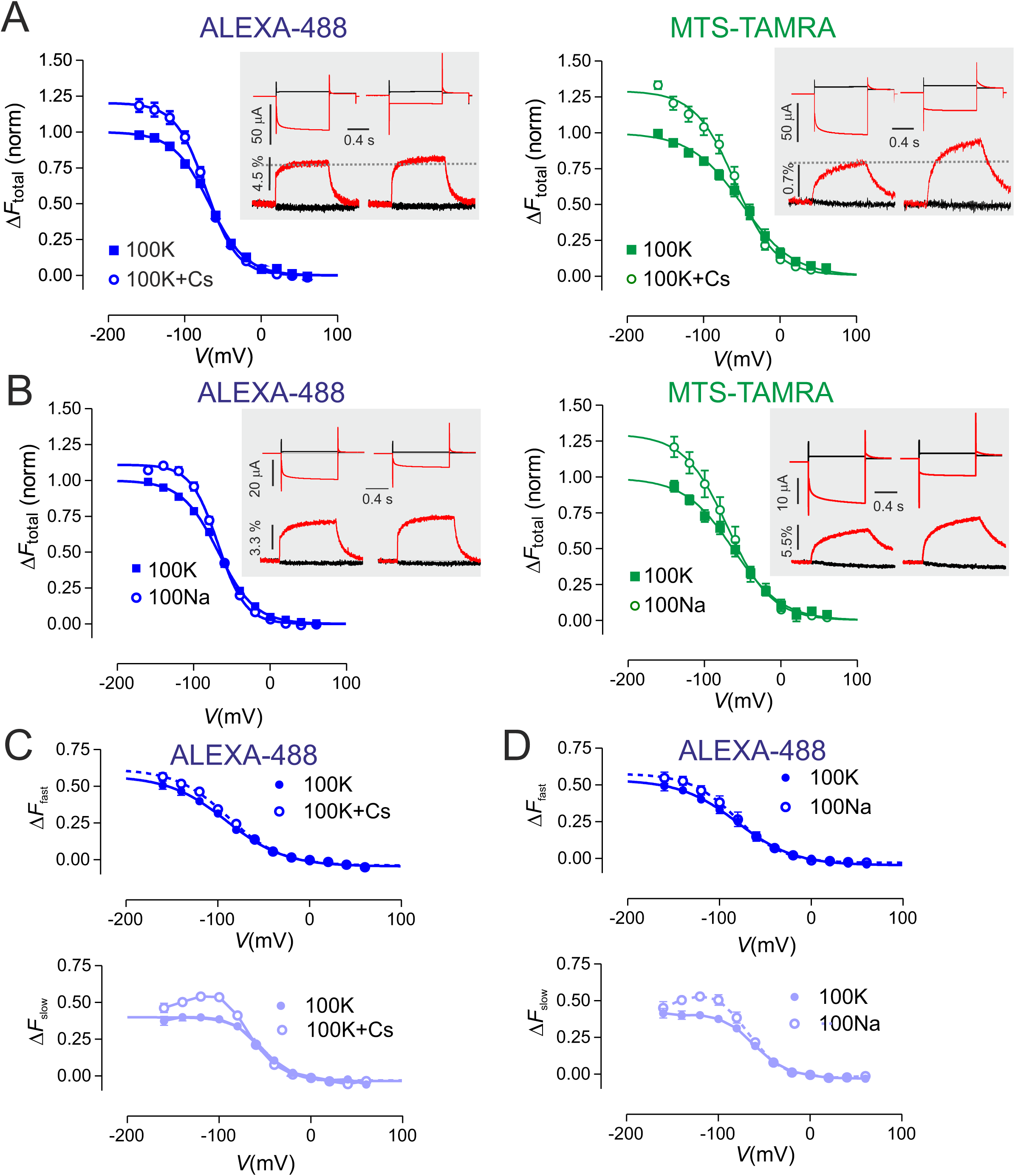
Δ*F* for both labels increases after changing permeation conditions. A) Δ*F-V* data before and after addition of 10 mM CsCl to bath containing standard 100K solution for spHCN^R332C^ expressing oocytes labeled with ALEXA-488 (left panel) and MTS-TAMRA (right panel). Insets show representative membrane current (upper traces) and Δ*F* (lower traces) for voltage steps to -160 mV and +20 mV from 0 mV holding potential and Δ*F* shown as % of background fluorescence. In both cases, the total Δ*F* was measured at the end of the test pulse. Δ*F-V* data were fit with the Boltzmann equation (Eqn 1) (continuous lines). Parameters given in Table 2. Δ*F-V* data were normalised to control condition (100K superfusion) for n= 9 replicates. B) Δ*F-V* data before and after replacing bath solution with 100Na solution to give 2 mM KCl for spHCN^R332C^ expressing oocytes labeled with ALEXA-488 (left panel) and MTS-TAMRA (right panel). Insets show representative membrane current (upper traces) and Δ*F* (lower traces) for voltage steps to -160 mV and +20 mV from 0 mV holding potential and Δ*F* shown as % of background fluorescence. Data analysis as in A) for n= 5 replicates. C) Δ*F-V* data for ALEXA-488 labeling showing fast (upper panel) and slow (lower panel) components before and after addition of 10 mM CsCl to bath as in A). Data were analysed by fitting Δ*F_total_* with a single exponential plus variable offset (see Materials and Methods) and normalised with respect to Δ*F_total_^max^*. D) D*F-V* data for ALEXA-488 labeling showing fast and slow components before and after replacement of 100K with 100Na solution to give 2 mM KCl as in B). Data were analysed by fitting Δ*F_total_* with a single exponential plus variable offset (see Materials and Methods) and normalised with respect to Δ*F_total_^max^*.

**Table 2.** Comparison of Boltzmann fit parameters before and after changing ion composition in pore.

a) 100K solution supplemented with 10 mM CsCl
|  | ALEXA-488 (11) |  | MTS-TAMRA (6) |  |
| --- | --- | --- | --- | --- |
|  | 100K | 100K+Cs | 100K | 100K+Cs |
| $\Delta F_{total}^{max}$ | 1.00±0.01 | 1.20±0.03 | 1.00±0.03 | 1.30±0.04 |
| $V_{0.5}^F$ (mV) | -67.8±1.1 | -72.2±1.6 | -49.7±3.4 | -58.2±2.9 |
| $z^F$ | 1.09±0.05 | 1.27±0.08 | 0.77±0.06 | 0.96±0.08 |

b) Reducing external $K^+$ from 100 mM (100K solution) to 2 mM (100Na solution)
|  | ALEXA-488 (5) |  | MTS-TAMRA (4) |  |
| --- | --- | --- | --- | --- |
|  | 100K | 100Na | 100K | 100Na |
| $\Delta F_{total}^{max}$ | 1.00±0.01 | 1.10±0.02 | 0.99±0.06 | 1.30±0.09 |
| $V_{0.5}^F$ (mV) | -67.8±0.8 | -68.1±1.1 | -62.6±5.5 | -69.7±6.1 |
| $z^F$ | 1.08±0.04 | 1.44±0.08 | 0.81±0.09 | 0.90±0.12 |

c) Analysis of fast component of ALEXA-488 fluorescence after supplementing with 10 mM CsCl or $K^+$ reduction.
|  | ALEXA-488 (9) |  | ALEXA-488 (5) |  |
| --- | --- | --- | --- | --- |
|  | 100K | 100K+Cs | 100K | 100Na |
| $\Delta F_{fast}^{max}$ | 0.62±0.04 | 0.66±0.03 | 0.58±0.04 | 0.61±0.04 |
| $V_{0.5}^F$ (mV) | -90.2±4.5 | -88.5±3.3 | -80.5±4.6 | -81.5±4.4 |
| $z^F$ | 0.82±0.10 | 0.89±0.09 | 0.83±0.12 | 1.02±0.2 |

Values are shown as mean ±SE as reported by fit to pooled, normalized data using Boltzmann equation. Data for individual cells were normalised to control condition (100K) before pooling. Number of cells analysed given in parentheses. Boltzmann fit to $\Delta F_{fast}$ , normalized to $\Delta F_{total}^{max}$ . Shaded values after changing to 100K+Cs solution or 100Na solution as indicated.

To examine if the mode shift was reflected in Δ*F*, we used the prepulse protocol incrementing in 8 ms or 80 ms intervals with a deactivating step to +40 mV and assessed the behavior of tail currents and Δ*F* under three superfusion conditions as above. Fig 6A,C show representative data sets for ALEXA-488 and MTS-TAMRA labeling respectively, with 80 ms prepulse increments. For both labeling conditions, superfusion with 10 mM CsCl added to the external medium (100K+Cs) blocked the activating current and increased total Δ*F*. Moreover, the normalised tail currents relaxed with a longer delay (quantified as the time to half amplitude (*t*_0.5_*^tail^*) (see (Bruening-Wright and Larsson, 2007)) relative to the tail current delay at 80 ms (Fig 6B,D). When superfused with 100Na solution containing only 2 mM KCl, Δ*F* increased as for superfusion with 100K+Cs and *t*_0.5_*^tail^* was further delayed. We excluded tail data that was obviously contaminated by endogenous currents activated during longer prepulses, which most likely accounted for the greater variability between individual cells (see Discussion). We also confirmed that the dependence of *t*_0.5_*^tail^* on external superfusate was not an indirect effect of the labeling as qualitatively similar results were obtained with an unlabeled spHCN^R332C^ (Fig SI 1).

**Figure 6.**
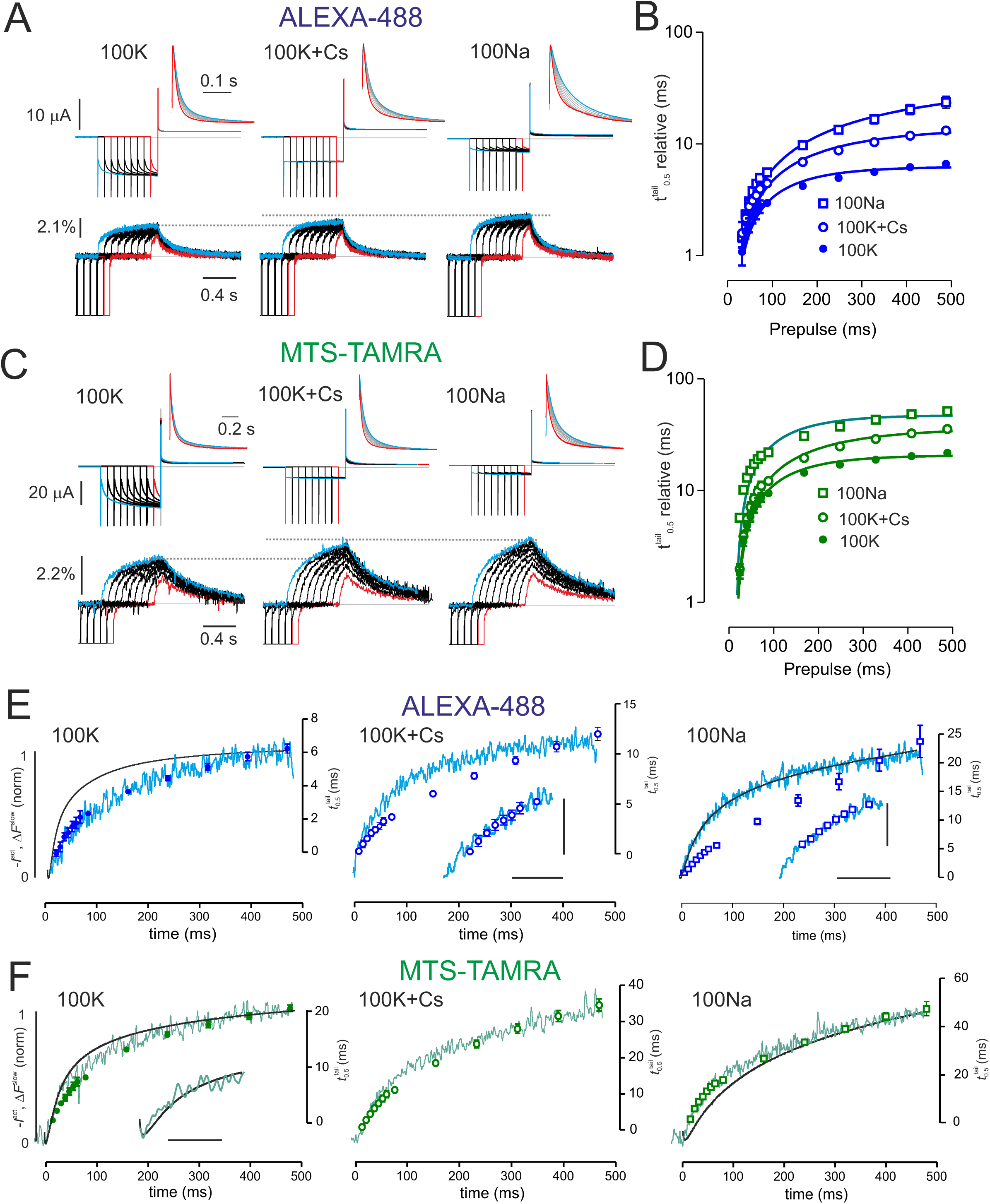
Fluorescence from spHCN^R332C^ channels labeled with ALEXA-488 and MTS-TAMRA report on mode shift development. A) Representative simultaneous current and Δ*F* recordings for an oocyte expressing spHCN^R332C^ labeled with ALEXA-488 and superfused in 100K, 100K+Cs and 100Na solutions in response to increasing prepulse steps to -160 mV, incremented by 80 ms from an initial 8 ms wide pulse. The traces are aligned at the prepulse OFF transition to +40 mV. Insets show respective normalized tail currents at +40 mV plotted on an enlarged time scale. To resolve the tail currents, capacitive transients were eliminated by subtracting the response to the 8 ms wide pulse to -160 mV. Δ*F* shown as % of background fluorescence. Dotted lines show that Δ*F* increases for superfusing with 100K+Cs and 100Na relative to 100K. The traces were signal averaged 4-fold to improve the signal-to-noise ratio. Δ*F* shown as a % of background fluorescence. B) Time for tail current to decay to 50% of its initial amplitude (*t*_0.5_*^tail^*) combining data with 8 and 80 ms pulse increments and shown relative to the tail response 16 ms pulse for three superfusion conditions plotted as a function of the prepulse width. Data shown as mean±SEM, n=10 (100K); 6 (100K+Cs); 6 (100Na). Points fitted with a single exponential function with time constants: 112± 12 ms (100K); 208±25 ms (100K+Cs) and 490±81 ms (100Na) (mean±SE). C) Representative simultaneous current and Δ*F* recordings for an oocyte expressing spHCN^R332C^ labeled with MTS-TAMRA, same conditions as in A). Insets show normalised tail currents as in A). Δ*F* shown as % of background fluorescence. Dotted lines show that Δ*F* increases for 100K+Cs and 100Na relative to the 100K condition. D) Time for tail current to decay to 50% of its initial amplitude (*t*_0.5_*^tail^*) combining data with 8 and 80 ms pulse increments as in B. Data shown as mean±SEM, n=9 (100K); 7(100K+Cs); 6(100Na). Points fitted with a single exponential function with time constants: 108 ± 7 ms (100K); 173 ± 15 ms (100K+Cs) and 105 ± 7 ms (100Na) (mean±SE). E) Correlations at -160 mV between *I^act^* (black trace) and the slow component of ALEXA-488 Δ*F* from same cell as in A (blue trace), for superfusion conditions as indicated. *I*^act^ is shown inverted, *I*^act^ and Δ*F* are normalised to corresponding values at 488 ms. Relative tail delay (*t*_0.5_*^tail^*) data from B) was overlaid on same abscissa according to the prepulse width so that the values at 16 ms and 488 ms match Δ*F* at these time points. Insets with Δ*F* trace expanded from 0-100 ms, show that for 100K+Cs and 100Na at short prepulse periods (up to 88 ms) there was an improved match between the time course of Δ*F* and *t*_0.5_*^tail^*. Scale bars for insets: vertical 5 ms, horizontal 50 ms. F) Correlations at -160 mV between *I^act^* (black trace) for the slow component of MTS-TAMRA Δ*F* (green trace) from same cell as in C, for superfusion conditions as indicated, plotted against time. *I*^act^ was inverted and Δ*F* normalised as in E). Inset for 100K superfusion shows expanded view to 88 ms obtained from same cell using the 8 ms time increment protocol with *I*^act^ and Δ*F* normalised to values at 88 ms time point. Scale bar = 40 ms. Relative tail delay (*t*_0.5_*^tail^*) data from D) overlaid on same abscissa according to the prepulse width according to the prepulse width so that the values at 16 ms and 488 ms match Δ*F* at these time points.

Fig 6E,F compares the time course of development of normalised *I*^act^ and Δ*F* with *t*_0.5_*^tail^* at −160 mV for prepulse widths up to 488 ms overlaid on the *I*^act^ and Δ*F* data, for each fluorophore. With ALEXA-488 labeling, there was no correlation between *I*^act^ and Δ*F* for 100K superfusion as initially observed (Fig 1 B), however Δ*F* tracked the time course of *t*_0.5_*^tail^* over the whole range examined. This supported the notion that under these conditions the slow component of ALEXA-488 fluorescence reported conformational changes associated with the development of the mode shift. However, for 100K+Cs and 100Na superfusion, *t*_0.5_*^tail^* only tracked Δ*F* for prepulse periods up to 100 ms (Fig 6E, insets), which indicated that under conditions of low availability of K^+^ ions in the pore, the slow component of ALEXA-488 also reported conformational changes associated with pore opening. For MTS-TAMRA labeling *I*^act^ and Δ*F* correlated for prepulse periods < 88 ms (Fig 6F, inset) as expected from the parametric plots (Fig 1B) and *t*_0.5_*^tail^* for prepulse periods > 88 ms tracked the time course of Δ*F*, consistent with MTS-TAMRA reporting both channel opening and mode shift conformational changes. For 100K+Cs superfusion Δ*F* and *t*_0.5_*^tail^* deviated in the early phase, whereas for 100Na superfusion Δ*F* and *t*_0.5_*^tail^* tracked over the whole of the time window investigated.

In summary, these findings provide new evidence that spHCN channels display mode shift in their deactivation kinetics that depends on the availability of K^+^ ions in the pore. The development of tail current delay with increasing hyperpolarization period at −160 mV can account for part of the Δ*F* reported by both labels at Cys332 during the channel activation phase.

### Quenching experiments and structural modelling suggest that the fluorophores report environments with different solvent accessibility during activation

To gain further insight into the underlying reasons for the different characteristics of each fluorophore we explored the effect of superfusion with the collisional quencher, iodide. Externally applied iodide should only cause fluorescent quenching of reactive sites if they are accessible to the external solution (Patti et al., 2016, Cha and Bezanilla, 1998, Mannuzzu et al., 1996). Addition of 50 mM I^-^ to the bath solution resulted in a reduction in background fluorescence by approximately 10% and 20% for ALEXA-488 labeled oocytes (n=9) (Fig 7A) and MTS-TAMRA labeled oocytes (n=8) (Fig 7C), respectively. Exposure to I^-^ had only a marginal effect on the steady-state open probability of the labeled channels and ̴10% increase in maximum conductance at −160 mV for both fluorophores (SI Fig 2). For these experiments K^+^ remained constant (100 mM) and we attributed this effect to changes of anion composition in the pore, which is known to alter the conductance of HCN channels (Wahl-Schott et al., 2005). For ALEXA-488 labeled oocytes, the predicted maximum total fluorescence (Δ*F_total_^max^*) at extreme hyperpolarizing potentials was reduced by ̴25%, however Δ*F* showed negligible change in *V*_0.5_^F^ or apparent valence (*z*^F^) as reported by a Boltzmann fit to the Δ*F-V* data (Fig 7B, Table 3). Moreover, in the presence of I^-^, both components of Δ*F* were reduced similarly without significant change in the midpoint voltage or apparent valence (Fig SI 2B). In contrast, for MTS-TAMRA labeled oocytes, a Boltzmann fit to Δ*F_total_* reported only a small reduction in Δ*F_total_^max^* accompanied by a 15 mV left shift in *V*_0.5_^F^ and reduced *z*^F^ (Fig 7D, Table 3). These findings indicated that during channel activation each fluorophore experienced a different I^-^-accessible environment as a consequence of voltage-induced conformational changes: for ALEXA-488 labeled oocytes, the reduction in Δ*F_total_* over the whole voltage range examined was consistent with this label remaining in a solvent accessible environment independent of its position, as it moved in response to S4 movement. In contrast, for MTS-TAMRA labeling, the fluorophore was less affected by I^-^ exposure, particularly at extreme hyperpolarizations, which suggested that it remained in a largely solvent inaccessible position.

**Figure 7.**
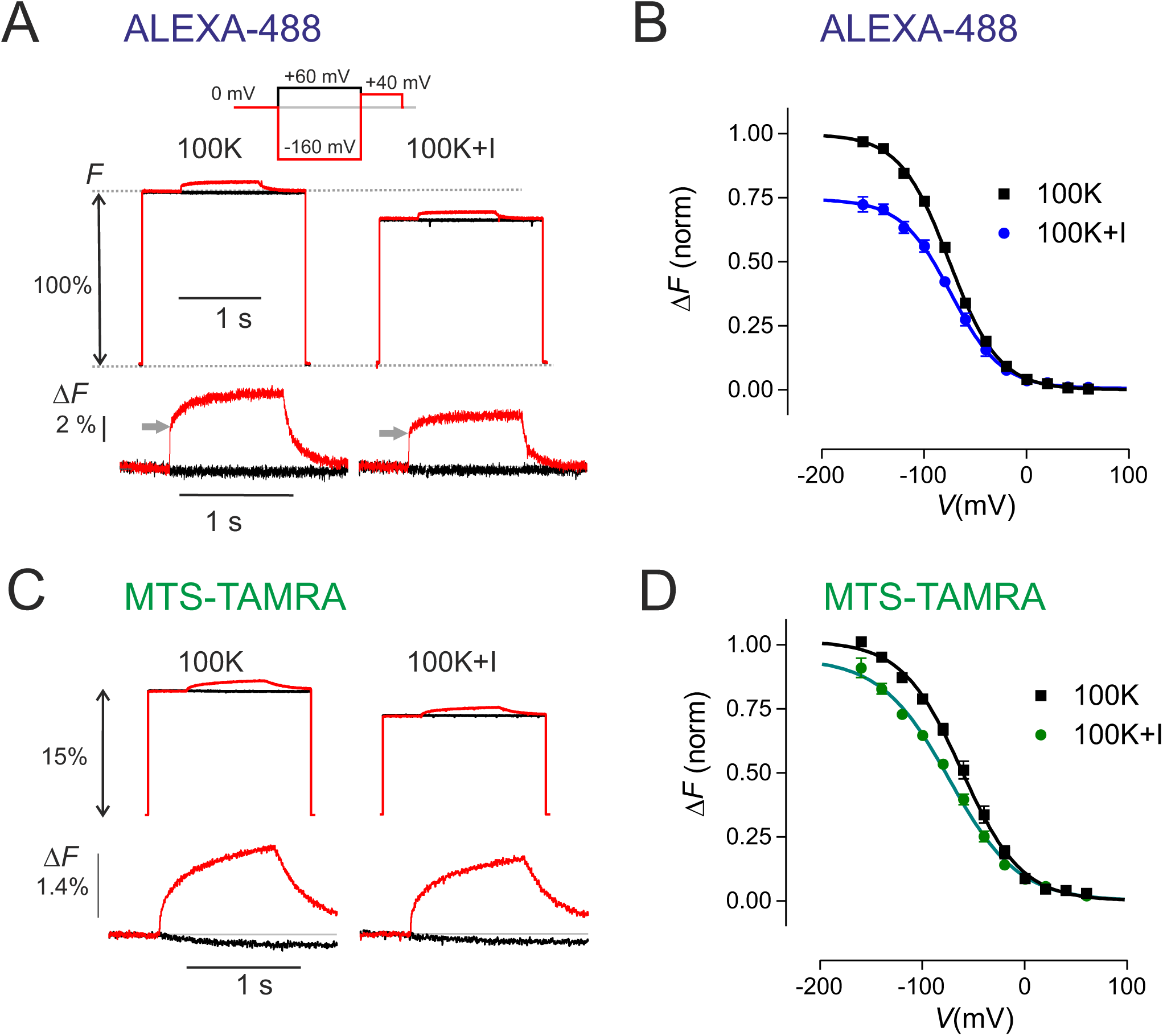
Iodide quenching experiments highlight differences in solvent accessibility for each fluorophore. A) Recordings of Δ*F* under control conditions (left traces) and following incubation in a solution for 30 s supplemented with 50 mM KI (see Materials and methods) for representative ALEXA-488 labeled oocyte. Upper traces: absolute fluorescence relative to the dark state; lower traces: an expanded view of Δ*F* in response to a voltage steps to +60 mV and -160 mV from 0 mV holding voltage. Arrows indicate Δ*F_fast_* component that is reduced after I^-^ exposure (see Fig SI 2). Δ*F* shown as % of background fluorescence. B) Δ*F_total_* plotted as a function of membrane potential pooled from (n=9) cells and fit with the Boltzmann equation. Data were normalised to the predicted Δ*F_total_^max^* from the fit to the control data set for each cell and pooled. The fit parameters are given in Table 3. C) Recordings of fluorescence under control conditions (left traces) and following incubation in a solution for 30 s supplemented with 50 mM KI (see Materials and methods) for a representative MTS-TAMRA labeled oocyte. Upper traces: absolute fluorescence relative to the dark state; lower traces: an expanded view of Δ*F* in response to a voltage steps to +60 mV and -160 mV from 0 mV holding voltage. Δ*F* shown as % of background fluorescence. D) Δ*F_total_* plotted as a function of membrane potential pooled from (n=11) cells and fit with the Boltzmann equation. Data were normalised to the predicted Δ*F_total_^max^* from the fit to the control data set for each cell and pooled. The fit parameters are given in Table 3.

**Table 3.** Comparison of Boltzmann fit parameters for spHCN^R332C^ fluorescence before and after exposure to 50 mM KI.

|  | ALEXA-488 (9) |  | MTS-TAMRA (11) |  |
| --- | --- | --- | --- | --- |
|  | 100K | 100K+KI | 100K | 100K+KI |
| $\Delta F_{total}^{max}$ | 1.00±0.02 | 0.74±0.03 | 1.01±0.02 | 0.95±0.03 |
| $V_{0.5}^F$ (mV) | -75.3±1.2 | -73.8±2.3 | -61.5±2.1 | -73.7±2.6 |
| $z^F$ | 1.06±0.05 | 1.05±0.09 | 0.89±0.05 | 0.77±0.04 |

Values shown as mean ±SE as reported by fit to pooled, normalised data using Boltzmann equation. Number of cells analysed given in parentheses. Data sets with iodide were normalised to $\Delta F_{total}^{max}$ for the control condition for each cell before pooling. Shaded values after iodide exposure.

To examine if these experimental findings had any structural correlates, we predicted binding poses for each fluorophore covalently bound to Cys332 for the closed (depolarized) and open (hyperpolarized) structures as shown in Fig. 8. For MTS-TAMRA labeling, under both depolarizing (Fig. 8A, left panel) and hyperpolarizing (Fig. 8A, right panel) conditions, the fluorophore adopted a ‘downward’ facing orientation, which placed the bulk of the molecule deep within the bilayer core. Although the immediate environment remained similar under both conformations, the identities of residues surrounding the fluorophore changed markedly (Fig. 8A, insets). In the depolarized (closed) state, 2-D ligand interaction diagrams predicted that the rhodamine head-group of MTS-TAMRA responsible for fluorescence is surrounded by hydrophobic residues of S5 and S6. During hyperpolarization the pattern of surround residues changes to less polar ones thereby explaining reduced quenching.

**Figure 8.**
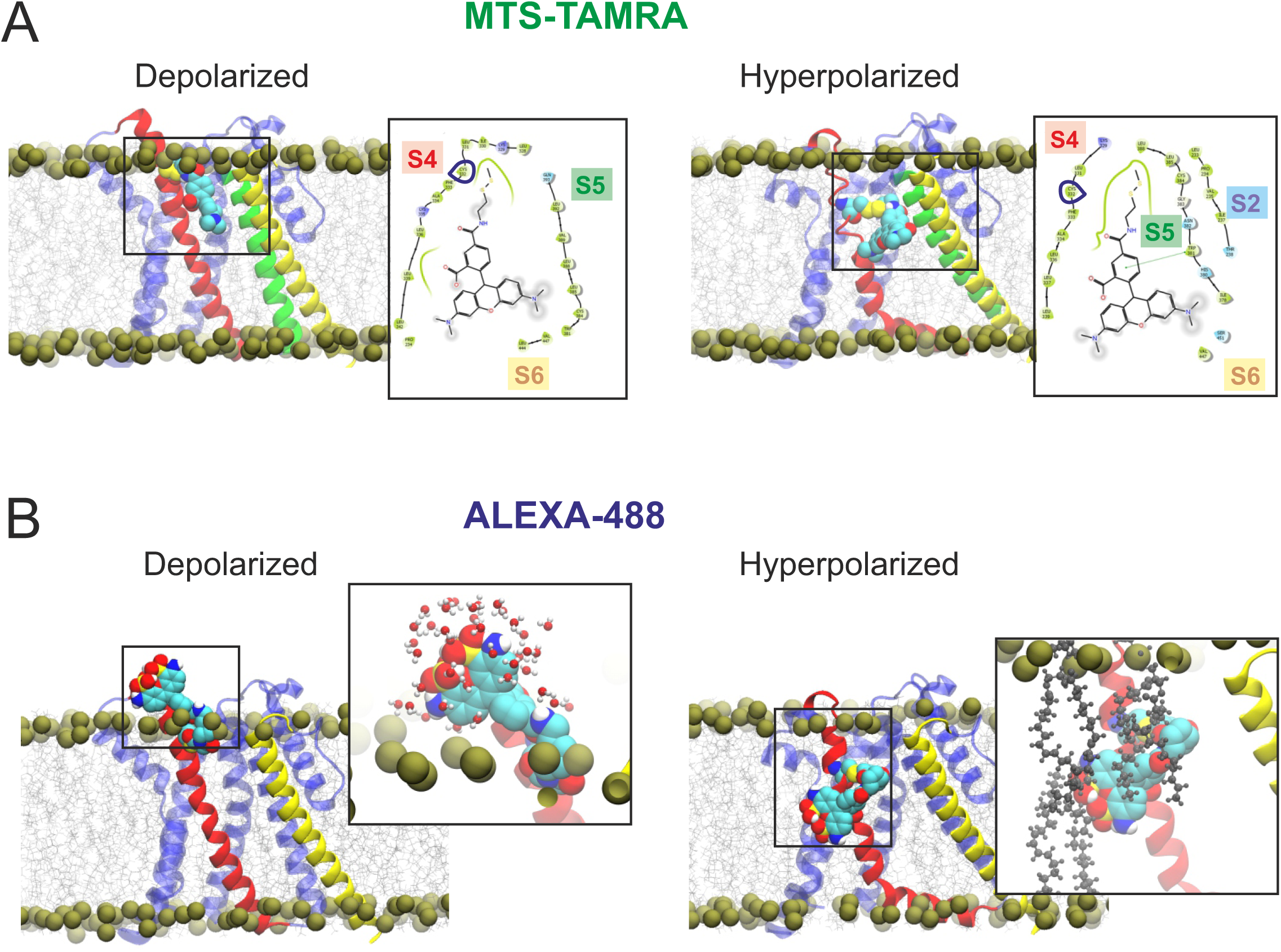
Molecular structures of the postulated binding poses predicted by covalent docking calculations. A) MTS-TAMRA docked at depolarized spHCN^R332C^ (left) and hyperpolarized spHCN^R332C^ (right), with insets showing predicted 2D ligand interaction diagrams. B) ALEXA-488 docked at depolarized spHCN^R332C^ (left) and hyperpolarized spHCN^R332C^ (right), with insets showing close-up rendering of the fluorophore interacting with either extracellular water (left) or lipid tails (right). One subunit of spHCN^R332C^ is represented by ribbons, with S4 in red, S5 in green and S6 in yellow. Fluorophores are shown as large brightly-colored spheres, bilayer phosphorus atoms are shown in bronze to illustrate the demarcation between the water and lipid phases. For the ligand interaction diagrams, residues in green are hydrophobic, residues in blue are polar, fluorophore atoms exposed to solvent shown by a cloud.

For ALEXA-488, (Fig 8B) there was a greater difference in its predicted orientation between the closed and open spHCN^R332C^ conformations. Under depolarizing conditions (Fig. 8B, left panel), one of the highest energetically favored orientations placed the bulk of the fluorophore in the extracellular space, allowing it to form a number of contacts with the top of the VSD, and thereby exhibit substantial solvent exposure. Under hyperpolarizing conditions, the fluorophore was predicted to re-orientate, with the bulk of the fluorophore now buried deep within the non-polar lipid tail region (Fig. 8B, right panel and inset). The marked difference in orientation between the two states reflects the significant voltage-dependent movements of S4, which would result in the fluorophore crossing through the juxtamembrane interface at hyperpolarizing potentials corresponding to the downward S4 helix movement. These movements would lead to a reduction of polarity in the microenvironment surrounding the fluorophore. Furthermore, ALEXA-488 was predicted to form contacts mainly with S4 itself, rather than S6, in contrast to MTS-TAMRA and consistent with ALEXA-488 reporting changes in its microenvironment closely associated with the initial rapid VSD movement.

## Discussion

We have investigated the changes in fluorescence (Δ*F*) recorded from labeled spHCN channels in response to voltage steps using two fluorophores (ALEXA-488, MTS-TAMRA) separately linked to the same site (Cys332) in the VSD S4 helix. Our findings using ALEXA-488 corroborate and extend those from an earlier study (Bruening-Wright et al., 2007). In brief, we found that ALEXA-488 reported Δ*F* consistent with the proposed initial rapid downward movement of S4 that precedes channel opening, together with subsequent conformational changes associated with channel opening and a mode shift that confers hysteresis to the channel open probability. In contrast, MTS-TAMRA reported conformational changes associated with the early phase of channel opening and mode shift. Thus, despite being covalently linked to the same site, distinct aspects of S4 movement were reported by each label during channel activation.

### ALEXA-488 reports on fast and slow VSD movement; MTS-TAMRA reports on slow VSD movement

Our data for ALEXA-488 labeling of Cys332 are in general agreement with those of Bruening-Wright and colleagues, whereby Δ*F* comprises two resolvable components (Bruening-Wright et al., 2007). They concluded that Δ*F* from ALEXA-488 labeled spHCN^R332C^ channels represented two distinct conformational changes of S4 that precede channel opening. Δ*F* from other labeled sites in S4 located closer to the N-terminal end of S4 and extracellular milieu (328-331) were similarly described by two exponential components, whereas for even more proximal sites (324–327), Δ*F* was best described by a single exponential (Bruening-Wright et al., 2007, Bruening-Wright and Larsson, 2007). Furthermore, Δ*F* correlated strongly with gating charge movement induced by voltage steps, which established that it reflected movement of S4 in response to the direction of the transmembrane electric field. The general consensus from these studies is that the faster component of Δ*F* represented S4 movement prior to channel opening, whereas the slower component reported the development of the mode shift following channel activation (Bruening-Wright et al., 2007, Elinder et al., 2006, Mannikko et al., 2002). Although our data appear qualitatively similar, there were differences when comparing the absolute values for time constants reported by the respective fit procedures. These might arise because the double exponential curve fitting in the earlier study was performed over a shorter time window (0.3 s, compared with 1 s in the present study) and we used single exponential fitting procedures to resolve the components with two time windows and signal averaging was employed for best resolution of the fast component.

Importantly, we now report that after block of channels with Cs^+^ or a 50-fold reduction in external K^+^ (100Na solution), the voltage dependence of the fast component was only marginally affected. This was consistent with the notion that this component of S4 movement occurred independently of subsequent VSD-PD coupling initiated by the formation of a helix break in S4 (Wu et al., 2021). In contrast, the slow component of Δ*F* reported by ALEXA-488 was found to be sensitive to the cation composition in the pore but its estimated magnitude only deviated from the control superfusion (100K solution) for potentials at which channels began to open (i.e. below −60 mV). This behavior indicated that the microenvironment of ALEXA-488 also reported on conformational changes associated with VSD to PD coupling. This may occur, for example, through the establishment of new non-covalent linkages between S4 and S5 (Bruening-Wright and Larsson, 2007, Ramentol et al., 2020, Wu et al., 2021, Hung et al., 2021, Elbahnsi et al., 2022) following the initial inward movement of the VSDs.

A 12-state allosteric scheme has been proposed (Wu et al., 2021) in which the fast Δ*F* was associated with the rapid, independent activation of each of the four S4’s, followed by a second and concerted S4 movement that leads to channel opening at extreme hyperpolarization. In this model, states occupied after the mode shift transitions were not included. Experimental evidence for the second S4 step was obtained from fluorescence recordings using a double mutant spHCN^R332C/E356A^ in which a predicted hydrogen bond between E356 at the C-terminal end of S4 and N370 in S5 is disrupted. This effectively uncouples the fast S4 movement from channel opening. Δ*F* from this second component correlates with *G* but has the opposite sign of the fast component and it was suggested that it would be masked in the spHCN^R332C^ construct due to overlapping voltage dependencies (Wu et al., 2021). Our simulations using the same 12-state model, with parameters adjusted to fit our data, recapitulated the basic features of our Δ*F* recordings for each label, excluding the mode shift component (Fig 4) and reaffirm how masking of the second S4 step would arise for ALEXA-488 labeling. Taken together, these findings strongly suggest that ALEXA-488 Δ*F* reported an initial, rapid movement of the VSDs, together with a slower and complementary component that would correspond to the proposed concerted movement of S4 helices leading to channel opening. Thus, our simulations were able to account for the difference in Δ*F* obtained under the two labeling conditions. In contrast to ALEXA-488 that reports Δ*F* arising from both S4 steps, labeling with MTS-TAMRA reported Δ*F* that correlated well with the initial phase channel activation, corresponding to the proposed concerted movement of S4 subunits that immediately precede channel opening.

### Both ALEXA-488 and MTS-TAMRA labeling report on spHCN^R332C^ mode shift

We now show that both labels at Cys332 report a slow component of Δ*F* that correlated well with mode shift development. Mode shift, as evidenced by tail current delay relates to the amount of time channels are open and transition to a second mode having a different voltage dependency. This might arise in the case of HCN channels if under dynamic (non-equilibrium) conditions the energy involved in forming (activation) and subsequently breaking (deactivation) of non-covalent VSD-PD linkages is different, thereby manifesting as hysteretic behavior (Bruening-Wright and Larsson, 2007, Villalba-Galea and Chiem, 2020, Cowgill and Chanda, 2023).

In the presence of high (100 mM) K^+^ in the pore, the tail current delay time course closely tracked the slow component of Δ*F*, although the absolute times were different for each label. This reflected the effect of the different labels on the overall channel kinetics. Furthermore, we show that blocking channels by Cs^+^ or reducing the external K^+^ 50-fold, increased the tail current delay and a correlation with the slow component of fluorescence was only seen for prepulse periods < 100 ms. The dependence of tail current delay development on pore cation composition was not reported previously for spHCN channels (Bruening-Wright and Larsson, 2007, Elinder et al., 2006, Mannikko et al., 2005) and contrasts with the behavior found for mammalian HCN1 channels under comparable conditions (Mannikko et al., 2005). In that study, channel block by Cs^+^ or reducing the availability of K^+^ ions in the pore resulted in a significantly faster development of tail current delay (Mannikko et al., 2005), which is the opposite of what we report here for spHCN^R332C^ channels. This may relate to differences in the pore structure between mammalian and invertebrate isoforms that alter the binding affinity of K^+^ ions within the selectivity filter (Jackson et al., 2007, Mannikko et al., 2005).

With ALEXA-488 labeling and prepulse periods >100 ms, Δ*F* and *t*_0.5_*^tail^* deviated markedly and there was greater variation in the estimates for *t*_0.5_*^tail^* between individual oocytes. Although contamination from endogenous currents activated by the longer activation period cannot be excluded (Bruening-Wright and Larsson, 2007), the systematic dependence on prepulse period and good agreement between Δ*F* and *t*_0.5_*^tail^* with MTS-TAMRA labeling for times >100 ms under all three superfusion conditions, suggests another underlying cause. For example, it may reflect the unmasking of a negative component of Δ*F* in the presence of 100K+Cs or 100Na observed at large hyperpolarizations that would effectively reduce the mode-shift contribution to Δ*F_total_*. Recently Wu and colleagues identified a negative Δ*F* under similar experimental conditions to those in our study by using the spHCN^R332C/E356A^ channel to decouple the VSD and PD (Wu et al., 2021). Here, we have also documented a decrease of Δ*F_sloss_* with ALEXA-488 labeling at extreme hyperpolarizing potentials (Fig 5 C,D) that may correspond to the component identified by Wu and colleagues. Further investigation of the mode shift behavior at different potentials would be required to identify the origins of the discrepancy between Δ*F* and *t*_0.5_*^tail^*.

These experiments highlight that changes in availability of K^+^ ions in the pore significantly alter the mode shift kinetics and suggest that structural changes in the selectivity filter, which have been shown for KscA channels to alter their hysteretic properties (Tilegenova et al., 2017), also impact on the VSD movement in HCN channels. The detection of a cation-dependent mode shift in spHCN^R332C^ channels validates the use of this construct for further studies to elucidate hysteresis using the sea urchin HCN channel as a model system.

### Orientation of fluorophores and structural determinants of fluorescence dynamics

Covalently linking fluorophores to engineered cysteines for VCF studies is a well-established technique that has led to important insights into channel function (Priest and Bezanilla, 2015, Cowgill and Chanda, 2019). Nevertheless, some limitations are imposed on the interpretation of Δ*F* with respect to developing channel structure-function relationships and interpretations should take into account the following caveats. First, the substitution of the Cys itself can lead to altered channel kinetics, where removal of Arg332 led to a significant left shift in the voltage dependent probability of channel opening as previously reported (Mannikko et al., 2002). Second, labeling itself can alter the channel opening probability (Fig 2A). Indeed, MTS-TAMRA labeling was found to partially revert the opening probability of labeled spHCN^R332C^ channels towards that of spHCN^WT^. Given the bulkiness of the fluorophores and their intrinsic charges, it is not surprising that deviations from the characteristics of the unlabeled or native protein occur. For example, the positive partial charge of MTS-TAMRA might compensate the loss of charge of Arg 332, whereas ALEXA-488 has a negative partial charge due to the sulfonic groups. Moreover, the difference in tether length between ALEXA-488 and MTS-TAMRA with 5 and 3 carbon atoms respectively would also be expected to play a role by conferring more flexibility to ALEXA-488. In contrast, once covalently linked to Cys332, MTS-TAMRA would be expected to impose more steric constraints on the channel, as evidenced by the slower deactivation kinetics and mode shift development. Our structural model of MTS-TAMRA-linked spHCN^R332C^ suggest that MTS-TAMRA is in close direct contact with the pore-lining S6 helix, as well as a predicted cation-π interaction with Trp381 which may serve to stabilise the fluorophore’s association with the S5 helix responsible for transmission of conformational changes between VSD and PD. These close contacts with S5 and S6 may serve to introduce additional friction that affects conformational transitions associated with pore closure. On the other hand, ALEXA-488 forms few contacts with the S6 helix, and therefore exerts less direct interference with pore closure.

## Conclusions

Our study describes, for the first time, the use of MTS-TAMRA as a tool to investigate HCN channel biophysical properties using VCF. The findings complement and extend previous VCF studies that were based exclusively on ALEXA-488 labeling. We have established that MTS-TAMRA reports a delayed fluorescence component corresponding to early channel opening, in contrast to ALEXA-488 that reports the initial VSD movement and a concerted S4 movement that immediately precedes channel opening. Supporting evidence from structural modeling gives new insights into the underlying molecular basis for the observed fluorescence. Both probes report on the mode-shift of spHCN channels during activation that is also dependent on the pore cation composition. Mode-shift provides an important physiological mechanism for dynamically regulating rhythmicity in mammalian HCN channels, and our data highlights the use of the non-mammalian channel as a good model to investigate this biophysical property. In summary, our data provides novel structure-function insights into HCN channel dynamics by introducing and validating the use of MTS-TAMRA in VCF studies. It also highlights more generally that the use of different labels for VCF can probe distinct biophysical aspects of ion channel opening expanding the capabilities of this technique.

## Acknowledgements

We thank Dr Tim Karle (Florey Institute of Neuroscience and Mental Health) for advice on the VCF hardware and providing the excitation LEDs; Professor Ernest Wright and Dr Donald Loo (UCLA) for valuable comments on the manuscript and for technical discussions. This work was supported by the Victorian State Government Operational Infrastructure Support; the Australian Government National Health and Medical Research Council (NHMRC) independent research Institute Infrastructure Support Scheme (IRIISS); an NHMRC Program Grant (10915693) (to CAR) and the Research Training Group “Chemical biology of ion channels (Chembion)” funded by the German Research Foundation (DFG), Project number: 404595355, GRK: 2515 (to MNW). The spHCN^R332C^ construct was the kind gift of Prof. Peter Larsson (University of Miami) and spHCN^WT^ was the kind gift of Prof R Seifert (MPINB, Bonn). The authors declare no competing financial interests.

## Author Contributions

Conceptualization: ICF, CAR; Data curation: ICF, AH; Formal analysis: ICF, AH, MNW; Funding acquisition: CAR, MNW; Investigation: ICF, CEM, MNW; Methodology: ICF, AH; Project Administration: CAR, MSS; Resources: ICF, AK, CEM, MSS, MNW; Supervision: ICF; Writing original draft: ICF, MNW; Writing –revision/editing: ICF, AH, AK, CEM, CAR, MSS, MNW

---

**Figure SI1.** Mode shift as evidenced by tail current delay was also observed for unlabeled spHCN^R332C^. A) Normalised tail currents from a representative unlabeled cell expressing spHCN^R332C^ shown for three superfusion conditions as indicated using the voltage step protocol shown in inset with 80 ms increments in prepulse to -160 mV. B) Tail current delay, quantified as time to 50% of initial amplitude relative to 8 ms, plotted as a function of prepulse width for each superfusion condition.

**Figure SI2.**
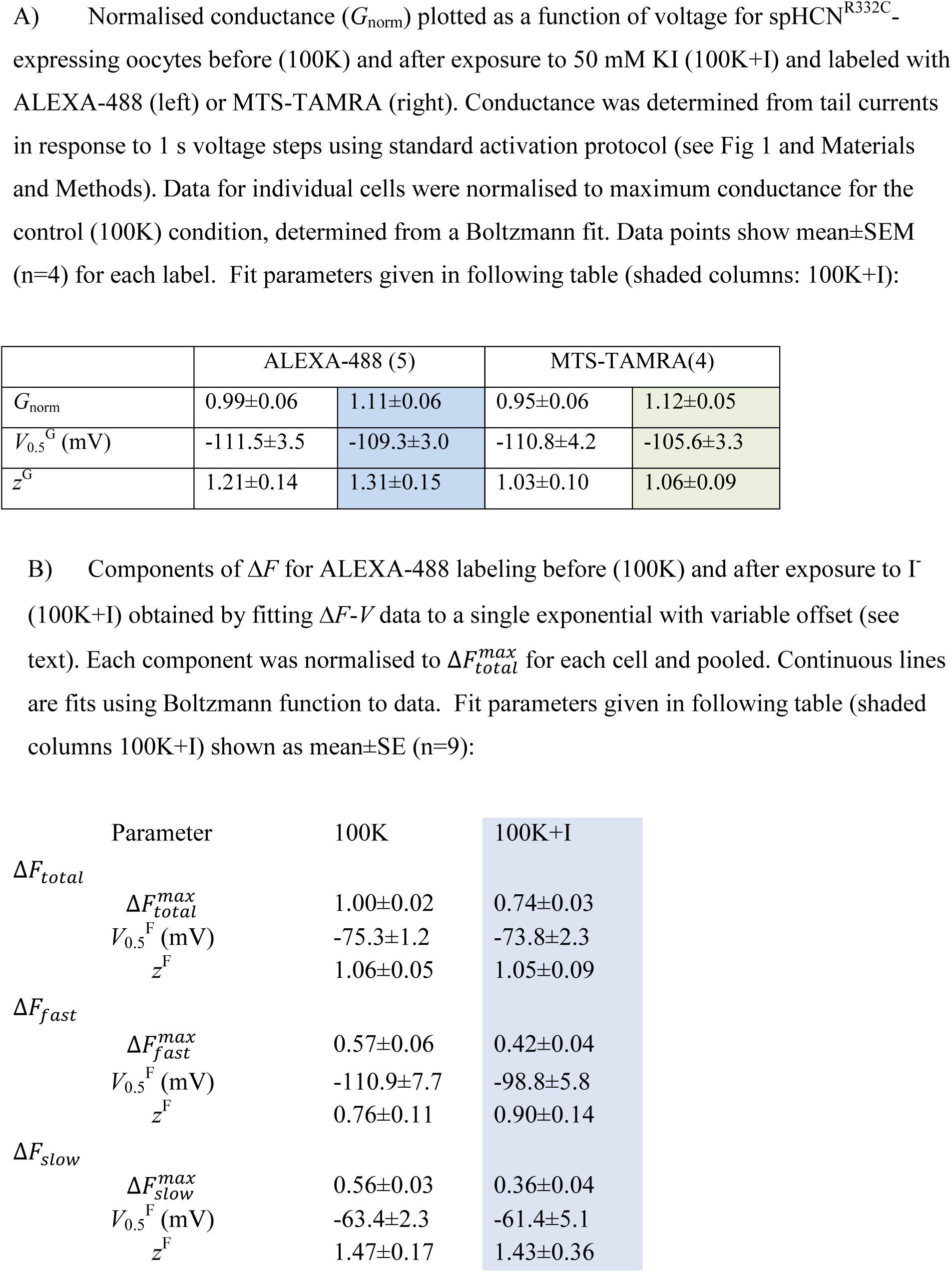
Normalised conductance increases during exposure to I^-^ without affecting voltage dependence.

**Table SI1.**
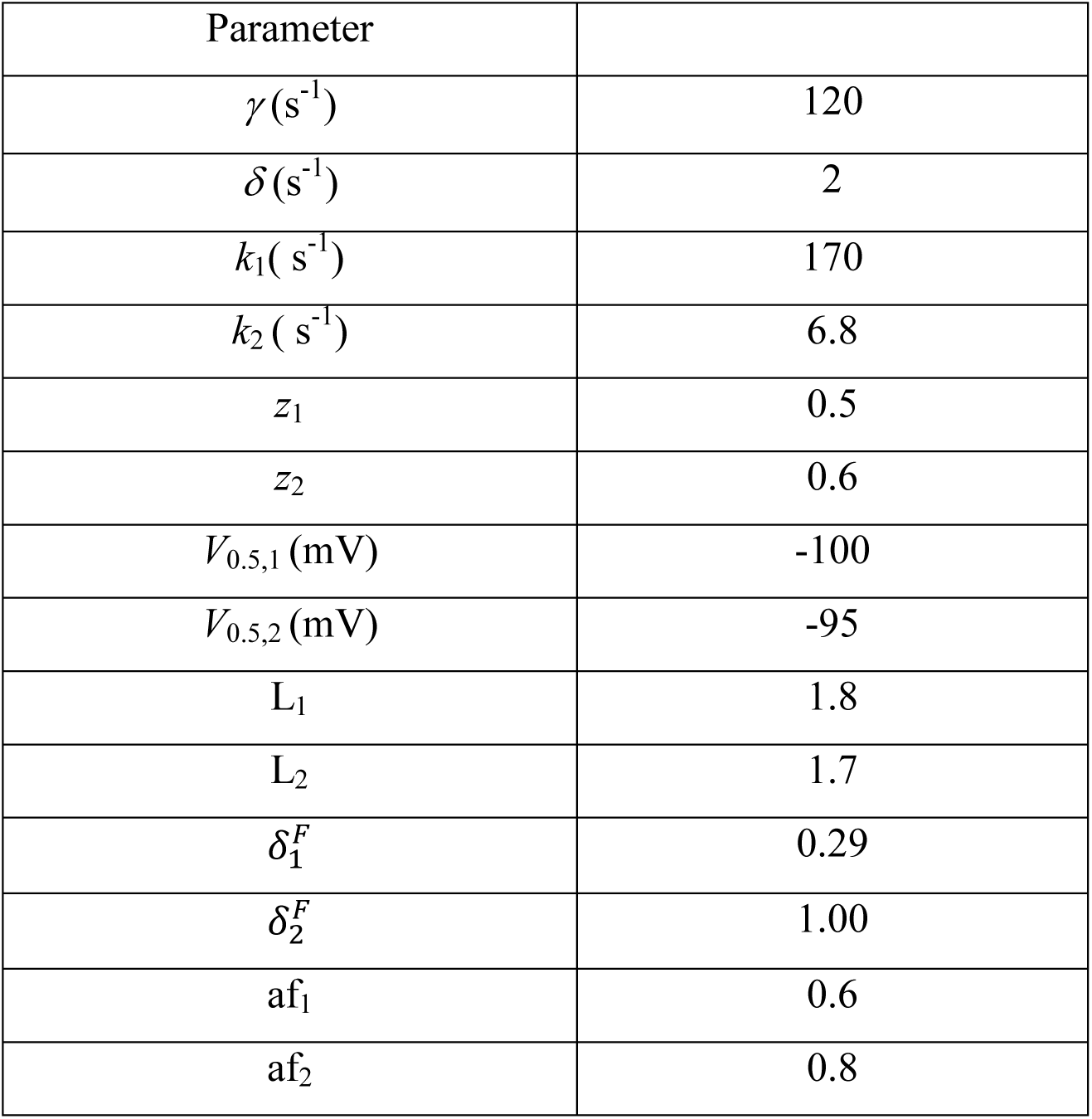
Simulation parameters. Parameters were determined by performing an initial fit using the inbuilt fitting routine in Madonna to a data set of activating currents (Fig 4B) and subsequently adjusted for the best fit by eye. The apparent fluorescence values (af_1,2_) are expressed in arbitrary units.

