## Supplemental text, tables for "Different fluorescent labels report distinct components of spHCN channel voltage sensor movement"

**Figure SI 1 Mode shift as evidenced by tail current delay was also observed for unlabeled spHCN$^{R332C}$**

A) Normalised tail currents from a representative unlabeled cell expressing spHCN$^{R332C}$ shown for three superfusion conditions as indicated using the voltage step protocol shown in inset with 80 ms increments in prepulse to -160 mV.

B) Tail current delay, quantified as time to 50% of initial amplitude relative to 8 ms, plotted as a function of prepulse width for each superfusion condition.

**Figure SI 2 Normalised conductance increases during exposure to I$^-$ without affecting voltage dependence.**

A) Normalised conductance ($G_{norm}$) plotted as a function of voltage for spHCN$^{R332C}$-expressing oocytes before (100K) and after exposure to 50 mM KI (100K+I) and labeled with ALEXA-488 (left) or MTS-TAMRA (right). Conductance was determined from tail currents in response to 1 s voltage steps using standard activation protocol (see Fig 1 and Materials and Methods). Data for individual cells were normalised to maximum conductance for the control (100K) condition, determined from a Boltzmann fit. Data points show mean±SEM (n=4) for each label. Fit parameters given in following table (shaded columns: 100K+I):

| | ALEXA-488 (5) | | MTS-TAMRA(4) | |
|---|---|---|---|---|
| $G_{norm}$ | 0.99±0.06 | 1.11±0.06 | 0.95±0.06 | 1.12±0.05 |
| $V_{0.5}^G$ (mV) | -111.5±3.5 | -109.3±3.0 | -110.8±4.2 | -105.6±3.3 |
| $z^G$ | 1.21±0.14 | 1.31±0.15 | 1.03±0.10 | 1.06±0.09 |

B) Components of $\Delta F$ for ALEXA-488 labeling before (100K) and after exposure to I$^-$ (100K+I) obtained by fitting $\Delta F$-$V$ data to a single exponential with variable offset (see text). Each component was normalised to $\Delta F_{total}^{max}$ for each cell and pooled. Continuous lines are fits using Boltzmann function to data. Fit parameters given in following table (shaded columns 100K+I) shown as mean±SE (n=9):

| | Parameter | 100K | 100K+I |
|---|---|---|---|
| $\Delta F_{total}$ | | | |
| | $\Delta F_{total}^{max}$ | 1.00±0.02 | 0.74±0.03 |

| | | | |
|---|---|---|---|
| | $V_{0.5}{}^{F}$ (mV) | -75.3±1.2 | -73.8±2.3 |
| | $z^{F}$ | 1.06±0.05 | 1.05±0.09 |
| $\Delta F_{fast}$ | | | |
| | $\Delta F_{fast}^{max}$ | 0.57±0.06 | 0.42±0.04 |
| | $V_{0.5}{}^{F}$ (mV) | -110.9±7.7 | -98.8±5.8 |
| | $z^{F}$ | 0.76±0.11 | 0.90±0.14 |
| $\Delta F_{slow}$ | | | |
| | $\Delta F_{slow}^{max}$ | 0.56±0.03 | 0.36±0.04 |
| | $V_{0.5}{}^{F}$ (mV) | -63.4±2.3 | -61.4±5.1 |
| | $z^{F}$ | 1.47±0.17 | 1.43±0.36 |

**Table SI 1.  Simulation parameters**

Parameters were determined by performing an initial fit using the inbuilt fitting routine in Madonna to a data set of activating currents (Fig 4B) and subsequently adjusting for the best fit by eye. The apparent fluorescence values are expressed in arbitrary units.

| Parameter | |
|---|---|
| $\gamma$ ($s^{-1}$) | 120 |
| $\delta$ ($s^{-1}$) | 2 |
| $k_1$( $s^{-1}$) | 170 |
| $k_2$ ( $s^{-1}$) | 6.8 |
| $z_1$ | 0.5 |
| $z_2$ | 0.6 |
| $V_{0.5,1}$ (mV) | -100 |
| $V_{0.5,2}$ (mV) | -95 |
| $L_1$ | 1.8 |
| $L_2$ | 1.7 |
| $\delta_1^F$ | 0.29 |
| $\delta_2^F$ | 1.00 |
| $af_1$ | 0.6 |
| $af_2$ | 0.8 |
