## Supplementary figures and images for "Different fluorescent labels report distinct components of spHCN channel voltage sensor movement"

### Supplemetal figs

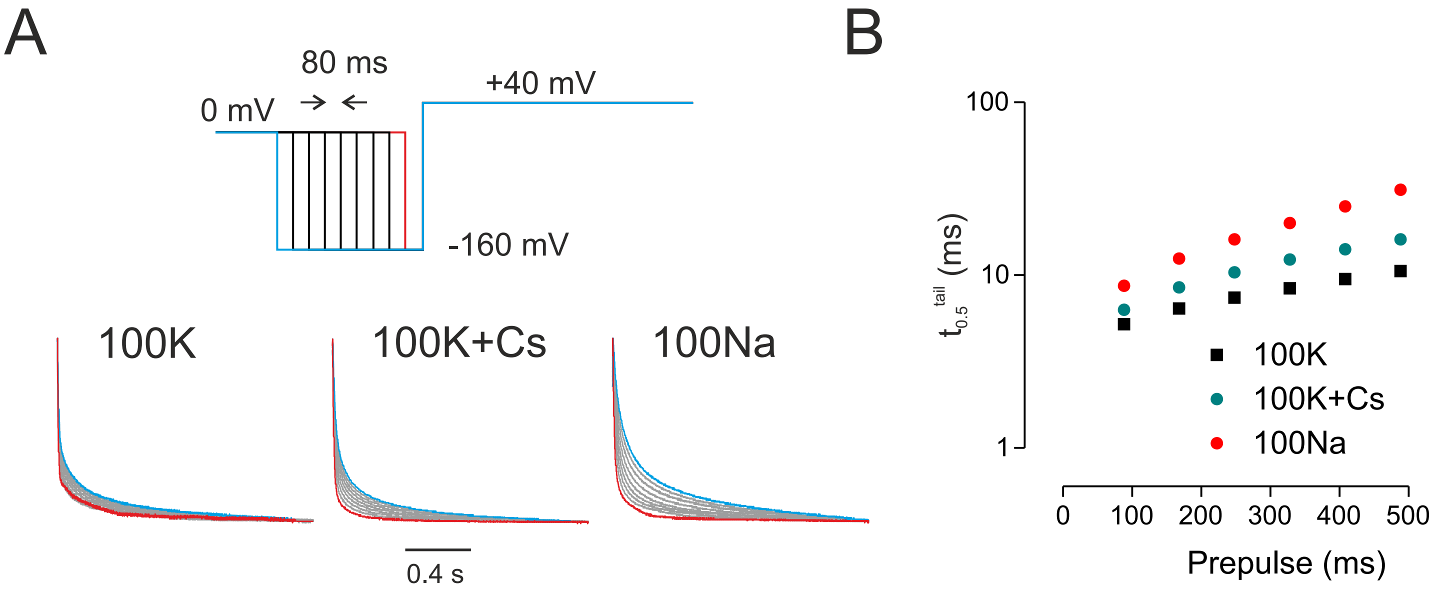

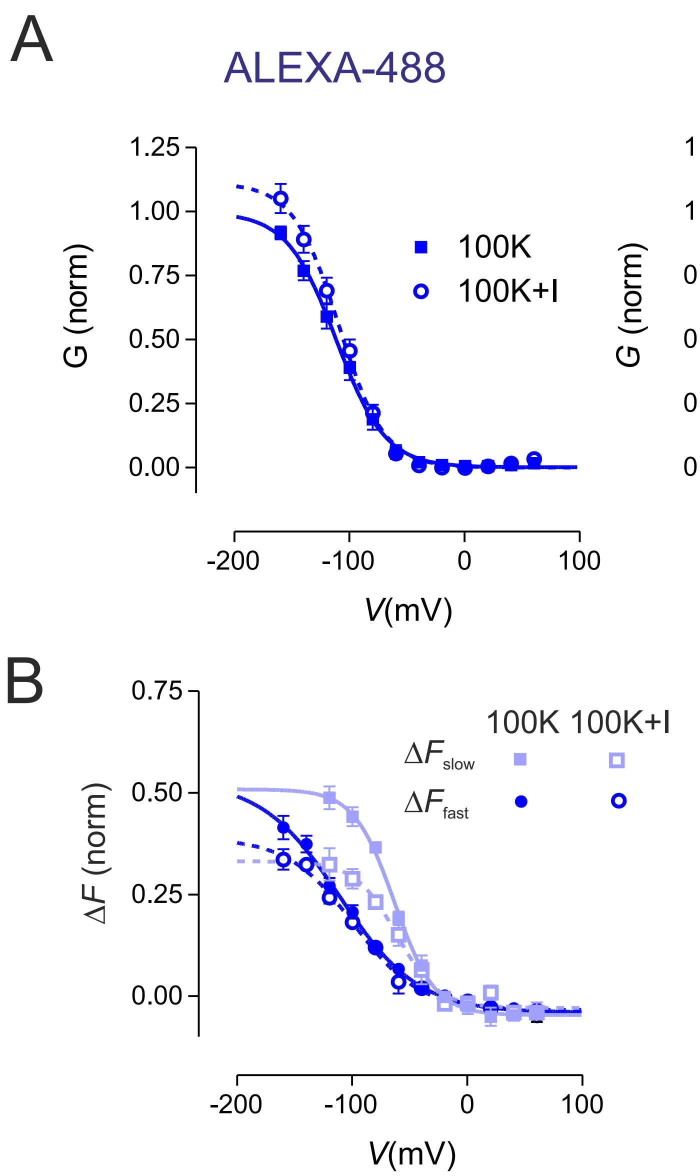
